# Non-shared dispersal networks with heterogeneity promote species coexistence in hierarchical competitive communities

**DOI:** 10.1101/2019.12.14.876383

**Authors:** Helin Zhang, Jinbao Liao

**Affiliations:** Ministry of Education’s Key Laboratory of Poyang Lake Wetland and Watershed Research, School of Geography and Environment, Jiangxi Normal University, Ziyang Road 99, 330022 Nanchang, China

**Keywords:** Dispersal heterogeneity, preemptive competition, competitive hierarchy, spatial coexistence, network theory, segregation-aggregation mechanism

## Abstract

The competition-colonization trade-off has been a classic paradigm to understand the maintenance of biodiversity in natural ecosystems. However, species-specific dispersal heterogeneities are not well integrated into our general understanding of how spatial coexistence emerges between competitors. Combining both network and metapopulation approaches, we construct a spatially explicit, patch-occupancy dynamic model for communities with hierarchically preemptive competition, to explore species coexistence in shared vs. non-shared dispersal networks with contrasting heterogeneities (including regular, random, exponential and scale-free networks). Our model shows that species with the same demography (i.e. identical colonization and extinction rates) cannot coexist stably in shared networks (i.e. the same dispersal pathways), regardless of dispersal heterogeneity. In contrast, increasing dispersal heterogeneity (even at very low levels of heterogeneity) in non-shared networks can greatly promote spatial coexistence, owing to the segregation-aggregation mechanism by which each species is restricted to self-organized clusters with a core of the most connected patches. However, these competitive patterns are largely mediated by species life-history attributes, for example, a unimodal biodiversity response to an increase of species dispersal rate emerges in non-shared heterogeneous networks, with species richness peaking at intermediate dispersal levels. Interestingly, increasing network size can foster species coexistence, leading to a monotonic increase in species-area curves. This strongly suggests that, unexpectedly, many more species can co-occur than the number of limiting resources. Overall, this modelling study, filling the gap between network structure and spatial competition, provides new insights into the coexistence mechanisms of spatial heterogeneity.

## Introduction

Exploring the mechanisms of community maintenance has been a central issue in ecology. Many mechanisms have been proposed (e.g. niche and neutral theories), and significant advances have been made in understanding species coexistence and consequently biodiversity maintenance (Chesson 2000; Hubbell 2001; Levine & HilleRisLambers 2009; Chu & Adler 2015). Among them, the competition-colonization trade-off has been a classic paradigm to explain the maintenance of biodiversity in natural ecosystems (Tilman 1994; Amarasekare 2000; Yu & Wilson 2001; Yu et al. 2004). But if there is no any tradeoff between species competition and demographic traits (colonization/extinction), how species can coexist stably in competitive communities has become a long-standing challenge for theoretical ecologists. Recently, non-hierarchical competition (i.e. competitive intransitivity) among species has been proposed as a potential endogenous mechanism for multispecies coexistence (Laird & Schamp 2006; Allesina & Levine 2011; Soliveres et al. 2015; Levine et al. 2017). However, a key question remains unsolved in hierarchical (transitive) competitive systems proposed by Tilman et al. (1994): whether there exists any other factor fostering species coexistence in such system without involving the colonization-competition trade-off. To probe the possible underlying mechanism of such coexistence, we shift our focus to species dispersal in spatially patchy environments. Most previous models of spatial coexistence assumed that species dispersal is lattice- or randomly-structured in two-dimensional space, ignoring the heterogeneity of dispersal networks (i.e. variation in patch connectivities) across the landscape (but Hanski & Ovaskainen 2000).

In nature, evidence of heterogeneous dispersal networks abounds (Urban & Keitt 2001; Fortuna et al. 2006; Grilli et al. 2015). For example, Kininmonth et al. (2010) found that the 321 reefs of the Great Barrier Reef constitute a scale-free small-world dispersal network for the species, where most reefs have only one or a few links and a very small proportion of reefs are extremely well-connected, following a power-law degree distribution. Montoya et al. (2008) observed that trees with bird-dispersed seeds perceive the landscape as a heterogeneous network, as opposed to trees with wind-dispersed seeds which move through a homogeneous network. Fortuna et al. (2006) identified a large spatial network of temporary ponds as breeding sites for amphibian species, observing a shift from a power-law to a truncated power-law distribution as the level of drought increases. As such, there has been an increasing interest in characterizing the persistence and dynamics of interacting species with dispersal heterogeneity using network theory (Urban & Keitt 2001; Bode et al. 2008; Holland & Hastings 2008; Dale & Fortin 2010; Kininmonth et al. 2010; Gilarranz & Bascompte 2012; Grilli et al. 2015; Gilarranz et al. 2017). In these representations, each network is described as a graph consisting of a set of nodes (i.e. colony sites or patches), and links between these nodes indicate dispersal pathways of individuals or sub-populations (Fortuna et al. 2006, 2009). These studies found that spatial heterogeneous networks greatly promote species persistence relative to the homogeneous networks by increasing local recolonization opportunities, demonstrating the importance of dispersal network structure for ecological dynamics (e.g. Holland & Hastings 2008; Gilarranz & Bascompte 2012).

Despite these advances, species-specific dispersal network connectivities have not been well integrated into our general understanding of how coexistence emerges among species. Even though a few models have considered spatial dispersal with different heterogeneities, they still assumed that all species have the same dispersal pathways (i.e. shared dispersal networks; e.g. Holland & Hastings 2008). This assumption neglects the fact that different species may perceive the landscape differently and therefore display distinct dispersal pathways, shaping diverse patterns of patch connectivity (Bunn et al. 2000; Nicholson & Possingham 2006; Fortuna et al. 2009; Hirt et al. 2018). Furthermore, some species can exhibit markedly different dispersal patterns even though they disperse at similar times through similar mechanisms, for example, animal species have distinct movement pathways due to different feeding preference on different plant seeds (Nathan & Muller-Landau 2000; Kinlan & Gaines 2003; Becker et al. 2007; Clobert et al. 2009). In fact, such species-specific dispersal networks not only arise from niche differences (e.g. species difference in habitat preference), but also result from different vectors for dispersal among species (e.g. seed dispersal by animals, gravity or wind; Yeaton & Bond 1991; Monotoya et al. 2008; Germain et al. 2019) or different dispersal traits (e.g. short- or long-distance dispersal). Using the approach of allometric random walks to predict the links among patches, Hirt et al. (2018) found that animal movement speed strongly depends on body mass, and the degree of connectivity of a network follows an allometric relationship, with medium-sized animals covering longer distances. In addition, the locomotion mode (flying, running or swimming) can influence the species-specific network connectivity, with flying animals being able to connect more distant patches than running ones. Furthermore, Germain et al. (2019) applied satellite imagery to link dispersal modes to movement of dispersal vectors, and observed that hydrological networks, animal paths and distance can explain the richness of species dispersed by gravity, animals and wind, respectively. Thus, there is an urgent need for spatial coexistence theory to incorporate species-specific dispersal networks with heterogeneity that are widespread in nature (Amarasekare 2008).

In this study, we construct a spatially explicit patch-dynamic model for multiple species with hierarchically preemptive competition, in order to make a systematic comparative analysis of species coexistence in shared *vs*. non-shared dispersal networks with contrasting heterogeneities (defined as the extent of variation in patch linking degrees). The shared dispersal networks refer to networks with the same patches (or colony sites) and dispersal pathways, as opposed to the non-shared dispersal networks where a system consists of two or more networks with the same patches but different linking patterns (so-called multilayer networks; see Pilosof et al. 2017). In other words, the non-shared networks mean that each species has their own linking architectures over the landscape, independent of each other. Note that, it is not obligatory to require all species to have different characteristics of patch linking degree distribution (i.e. different dispersal heterogeneities) in the non-shared dispersal networks, i.e., species can have the same linking degree distribution (i.e. the same heterogeneity), but their overall dispersal pathways across the landscape should not be exactly the same (i.e. some links between patches are common for species). To characterize heterogeneity in patch connectivities, four typical types of spatial dispersal networks are considered (illustrated in Fig. 1): regular (Bascompte & Solé 1995), random (Erdös & Rényi 1959; Watts & Strogatz 1998), exponential and scale-free networks (Barabasi & Albert 1999). Using this model, we systematically explore: (i) whether and how competitors with dispersal heterogeneity can co-occur in shared vs. non-shared networks when they have the same demography; and (ii) which properties of dispersal network structure can best maintain species diversity.

**Figure 1.**
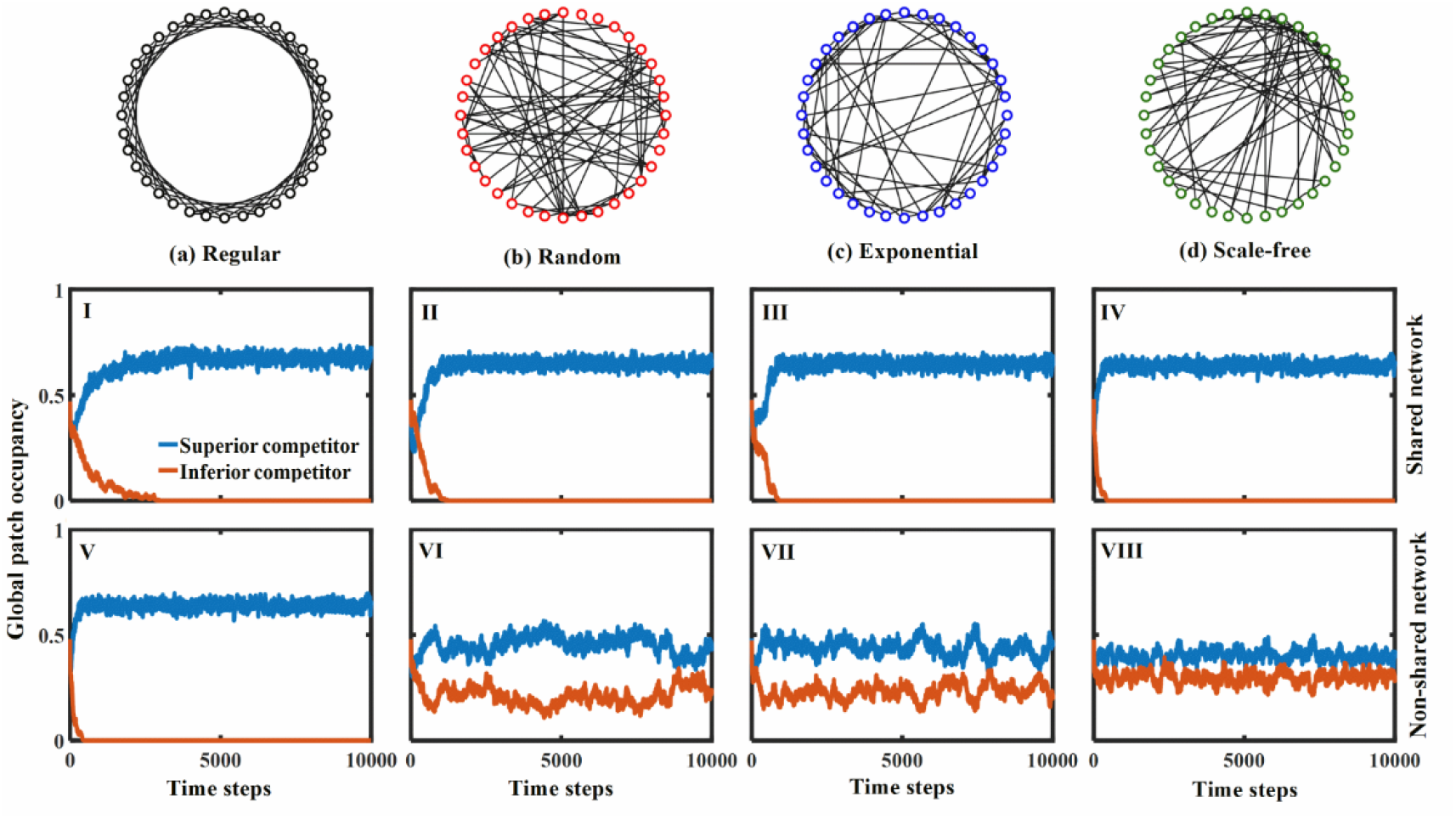
Patch dynamics of two competing species in shared (I-IV) vs. non-shared (V-VIII) dispersal networks, always containing 1024 patches and 2048 links with average linking degree 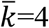. Four typical networks from the most homogeneous to the most heterogeneous are included: (a) regular, (b) random, (c) exponential and (d) scale-free networks. For display purposes, these networks consist of only 36 patches (nodes) with 72 links. In the shared networks, the dispersal pathways for both competing species are the same, while they are different in the non-shared networks but with the same heterogeneity. Parameter values are the same for both species: colonization rate *c*=0.05 and extinction rate *e*=0.05.

## Methods

### Dispersal networks with heterogeneity

We represent the landscape as a graph (spatial network) consisting of a set of nodes (patches or colony sites) connected by links. Each node denotes a patch linked with a number of other patches (i.e. linking degree *k*), and links between patches represent species dispersal pathways (i.e. functional connectivity among populations). As such, each type of dispersal network can be characterized by its linking degree distribution. Similar to Gilarranz & Bascompte (2012), four typical structures of dispersal networks are considered (illustrated in Fig. 1a-d):

i. A regular network where all patches have the same linking degree. For example, Figure 1a shows a completely homogeneous network where each patch has four links to other patches (*k*=4), equal to the lattice-structured model with nearest neighbour dispersal under periodic boundary conditions (Bascompte & Sole 1995; Hiebeler 2000; Liao et al. 2013a,b).
ii. A randomly structured network with randomly connected patches (Watts & Strogatz 1998). In particular, patch linking degrees follow a Poisson distribution with the variance equal to the average degree per patch (e.g. 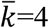 in Fig. 1b), thus there exists a small variation in patch linking degrees (Erdös & Rényi 1959).
iii. An exponential network constructed based on the generic algorithm of random attachment, i.e. the network continuously expands by the addition of new patches that are randomly connected to the patches already present in the system (see Barabási & Albert 1999). Figure 1c displays an exponential network with the same average linking degree 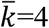 but more variability in linking degrees than the random network (i.e. higher heterogeneity with variance approximately equal to 5.8 in our model networks), following an exponential connectivity distribution (Fortuna et al. 2006).
iv. A scale-free network constructed according to the algorithm of Barabási & Albert (1999) with preferential attachment, leading to extremely high heterogeneity (with variance approximately equal to 27.1 in our model networks). This is a consequence of two generic mechanisms: (I) networks expand continuously by the addition of new patches, and (II) new patches attach preferentially to the patches that are already well-connected (Barabási & Albert 1999). Figure 1d exhibits such a network (keeping average linking degree at 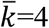), where most patches have only a few links, while a few patches are extremely well-connected, following a power-law degree distribution (Kininmonth et al. 2010).

In these networks, each patch is reachable for every species, that is, each patch has at least one link to another patch. Species are assumed to disperse equally in all directions with no preference, i.e. when patches *i*_1_ and *i*_2_ are linked, dispersal can occur from either *i*_1_ to *i*_2_ or vice versa.

### Competitive dynamics

We model a competition system structured in a large number of discrete habitat patches connected by species dispersal. Each patch can be vacant or host a single species. The system consists of *n* species with competitive hierarchy (i.e. ranking of species according to their competitive abilities), with the first species being the best competitor and the *n*-th species the worst (1>2>3>···>*n*). As colonizing a patch already occupied by another species may be intrinsically more difficult than colonizing an empty patch (e.g. plant propagule) because of the priority effect (Comins & Noble 1985; Calcagno et al. 2006; Fukami 2015), we mainly consider preemptive competition, i.e. species do not replace others but only compete for empty patches, with strong competitors having priority to colonize these patches. Thus, inferior species can colonize an empty patch only if superior species fail to establish there. All species in the landscape are assumed to have the same colonization (with rate *c*) and the same extinction (with rate *e*), in order to exclude any coexistence caused by the colonization-competition tradeoff (Tilman 1994). As such, we can unambiguously attribute any species coexistence to the explicit structural properties of our dispersal networks. In each time step, each occupied patch becomes extinct with a probability *e* regardless of species identity. At the same time, whether an empty patch is colonized depends on both species competitive hierarchy and abundances of its directly connected occupied patches. Therefore, the probability that a given empty patch *i* is colonized by the *S*-th competitor (1≤*S*≤*n*) should be

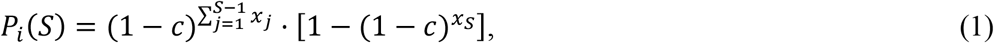

where *x_j_* (≥0) is the number of *j*-patches (occupied by species *j*) directly linked to the empty patch *i*, and 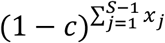 is the probability that patch *i* is unoccupied by those superior competitors (including species 1, 2, 3… *S*-1). Note that each empty patch is colonized by one of its directly connected occupied patches with an independent probability (*c*) regardless of species identity.

### Spatially explicit simulations

Initially each patch is occupied by a species randomly sampled from the species pool. For non-shared networks, we regenerate species-specific dispersal pathways according to their network properties (e.g. variation in patch linking degrees) and assign to each species randomly. Within each time step, we determine the local extinction for each occupied patch with a given probability (*e*). Then we calculate the probability that each empty patch becomes occupied by its directly connected species according to their competitive hierarchy as well as patch occupancies (see Eq. 1). Finally we record the patch occupancy for each species at each time step, calculated as their number of occupied patches divided by the network size (i.e. the total number of patches).

To determine the steady state, we preliminarily run the system for a long time, finding that 5000 time steps are sufficient to achieve stability. We eventually run each case until 10,000 time steps, and estimate patch occupancy for each species at steady state by averaging their occupancies between *t*=9000~10,000 time steps to avoid transient dynamics. Each case is explored with 100 replicates, starting from different dispersal architectures in each replicate but with identical network properties (e.g. the same network size, total links and their degree distribution). The mean of these 100 replicates (± standard deviation SD) yields species abundance at steady state. A broad range of biologically reasonable parameter combinations are explored and found to yield qualitatively similar outcomes (Figs S1-S21 in *Appendix*), thus allowing us to present our general results in Figs 1-5 by choosing one of those parameter combinations as a reference (cf. Gilarranz & Bascompte 2012; Liao et al. 2020). To reduce stochastic effects (Figs S1-S2 in *Appendix*), we model patch occupancy dynamics (via Maltab R2018b) using large networks consisting of 1024 patches and 2048 undirected links (cf. Gilarranz & Bascompte 2012). As such, all types of network have the same number of patches and links with the same average linking degree 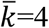, allowing us to compare species coexistence in dispersal networks with contrasting heterogeneities. Note that, in our model, we mainly focus on preemptive competition (i.e. species only compete for empty patches) without considering competitive displacement (i.e. the superior competitor can invade into and displace the inferior competitor; Tilman 1994), as we find that the assumption of competitive displacement is too strong to keep species coexistence if all species have the same demography (i.e. excluding the colonization-competition tradeoff) in hierarchical competitive communities (e.g. the inferior species would be always outcompeted; see Fig. S22 in *Appendix*).

**Figure 2.**
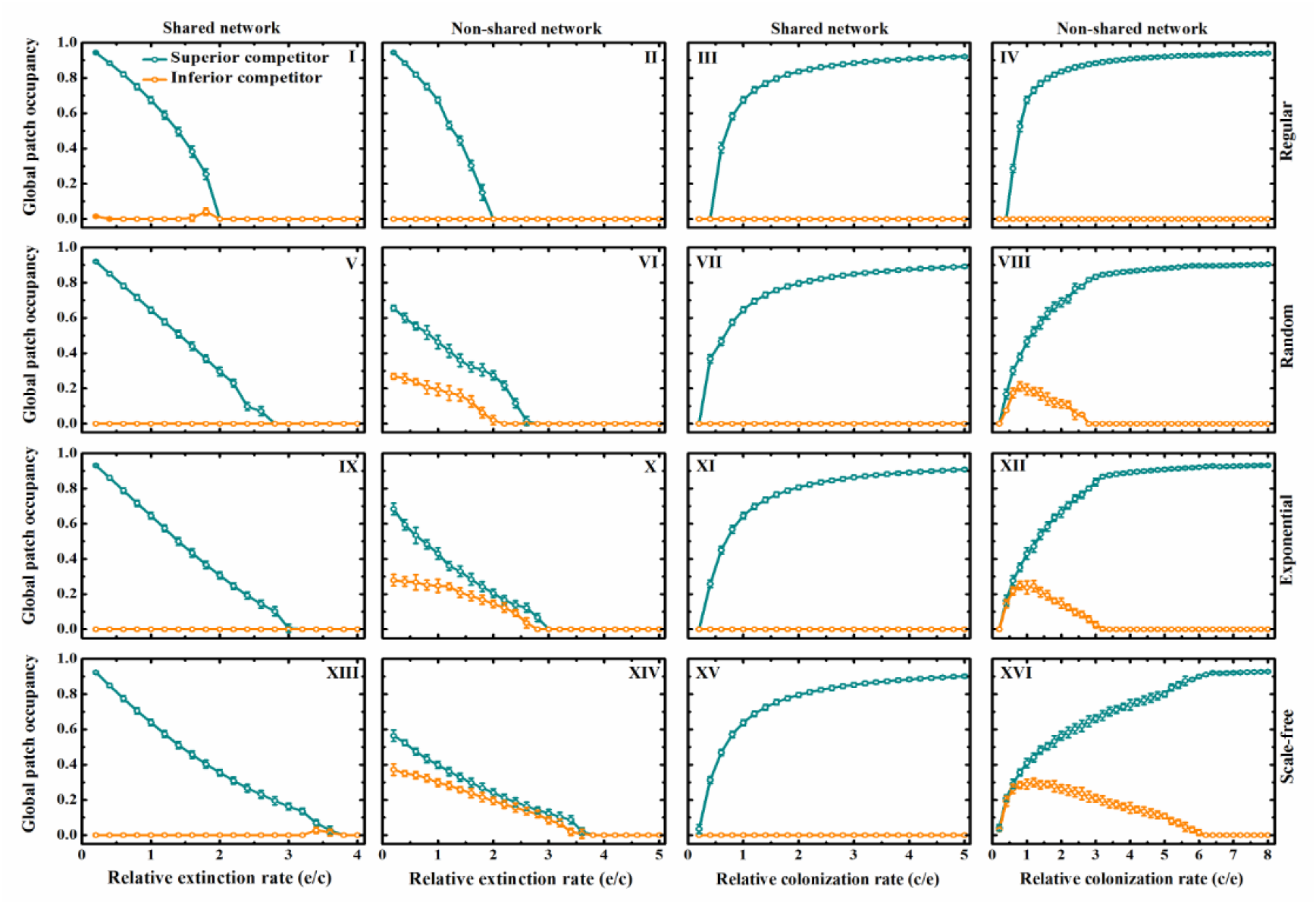
Effects of relative extinction (*e*/*c* at fixed *c*=0.05) and colonization (*c*/*e* at fixed *e*=0.05) rate on patch occupancy (mean ± standard deviation SD of 100 replicates) of both inferior and superior competitors at steady state in shared vs. non-shared networks with different heterogeneities, including regular, random, exponential and scale-free networks. These networks consist of 1024 patches and 2048 links, and non-shared networks indicate species-specific dispersal patterns for both competitors but with the same heterogeneity.

**Figure 3.**
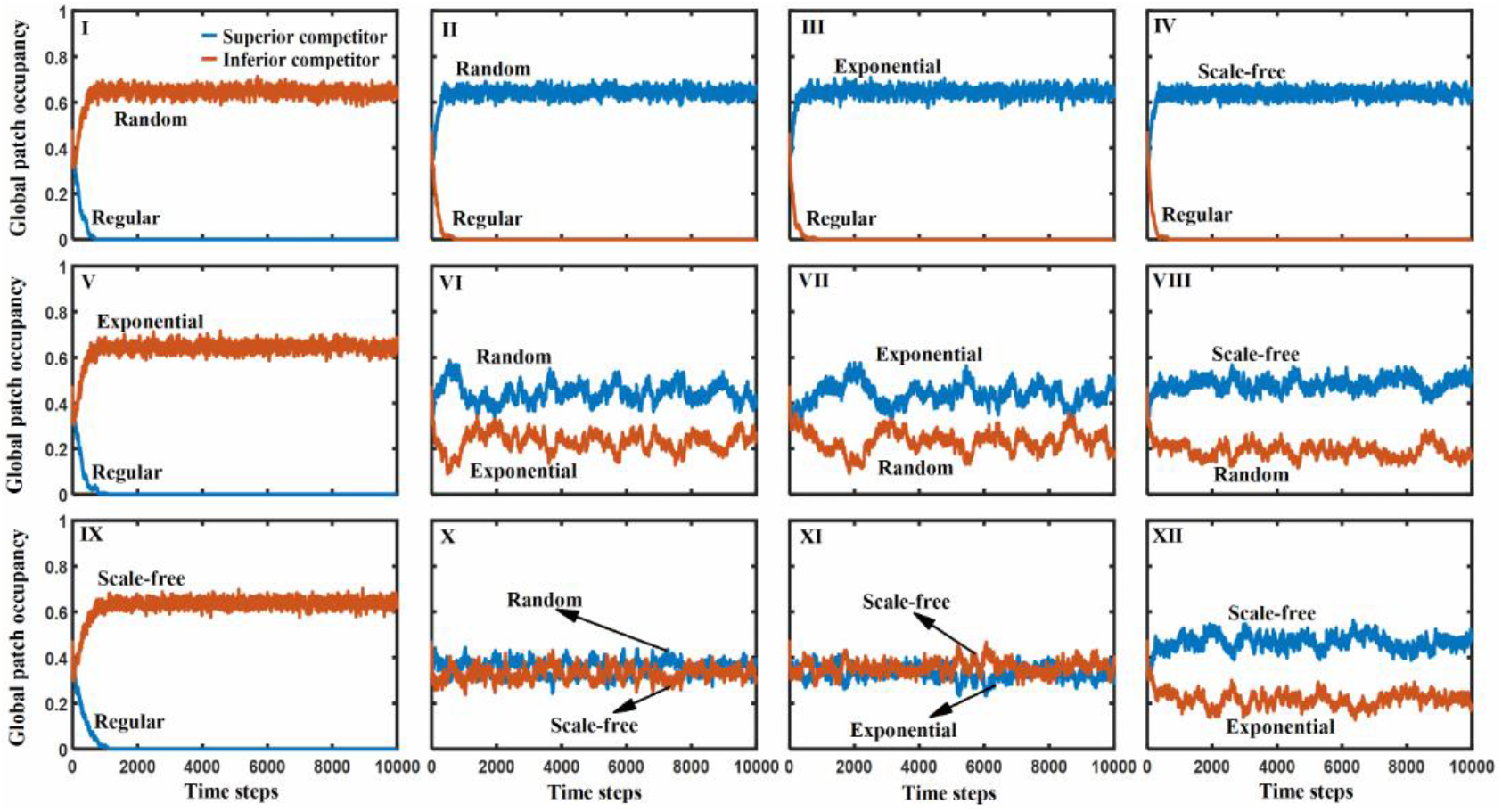
Patch dynamics of both inferior and superior competitors with different heterogeneous networks, consisting of 1024 patches and 2048 links. Four types of dispersal networks are considered: regular, random, exponential and scale-free networks. Parameter values for both species are the same: *c*=*e*=0.05.

**Figure 4.**
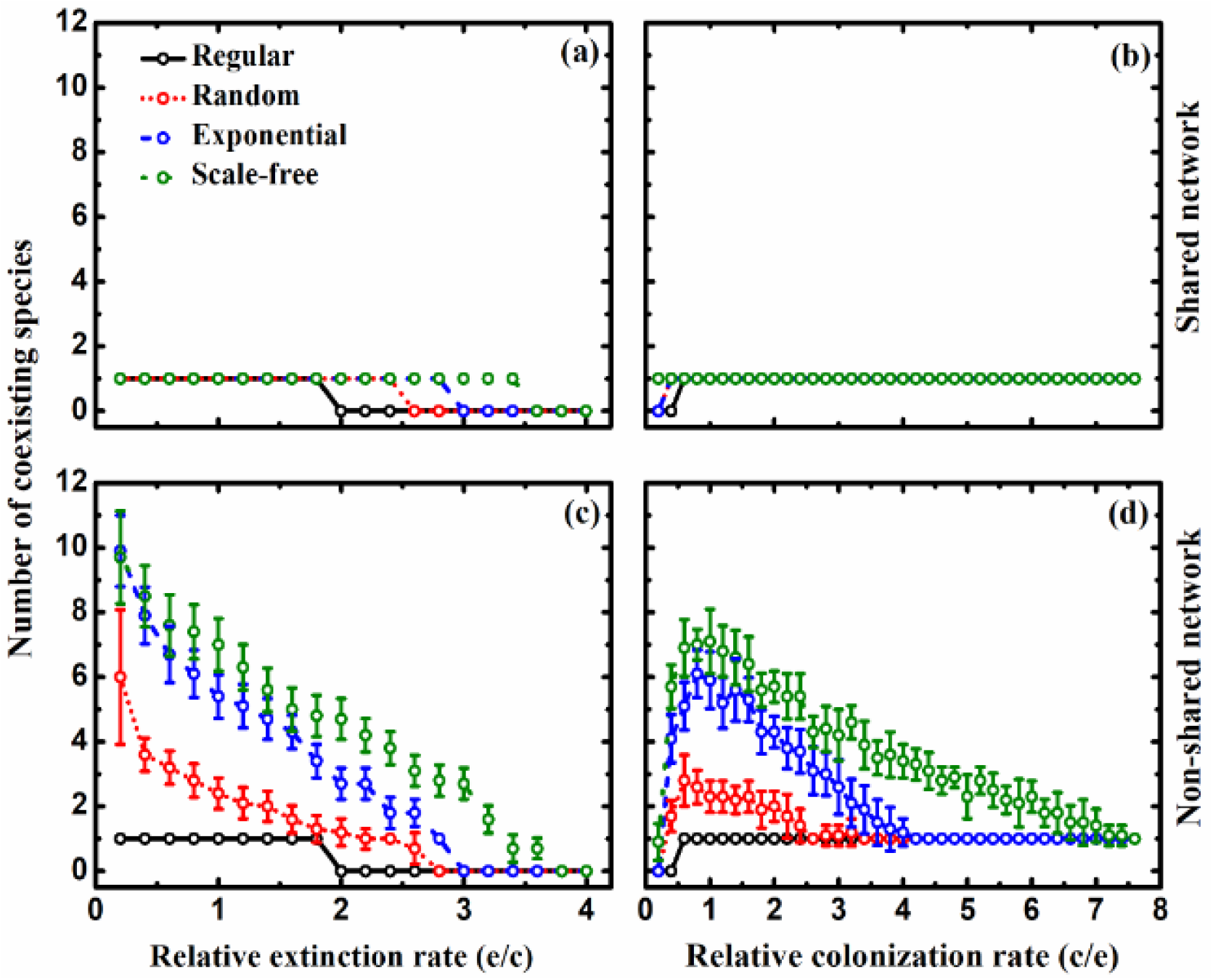
Effects of relative extinction (*e*/*c* at fixed *c*=0.05) and colonization rate (*c*/*e* at fixed *e*=0.05) on the number of coexisting species at steady state (mean ± SD of 100 replicates) in hierarchical competitive communities with preemptive competition, where species compete only for empty patches. All species are assumed to have the same colonization and extinction rates. Networks are either shared (graphs a & b) or non-shared but with the same heterogeneity (graphs c & d), and include regular, random, exponential and scale-free networks (again with 1024 patches and 2048 links).

**Figure 5.**
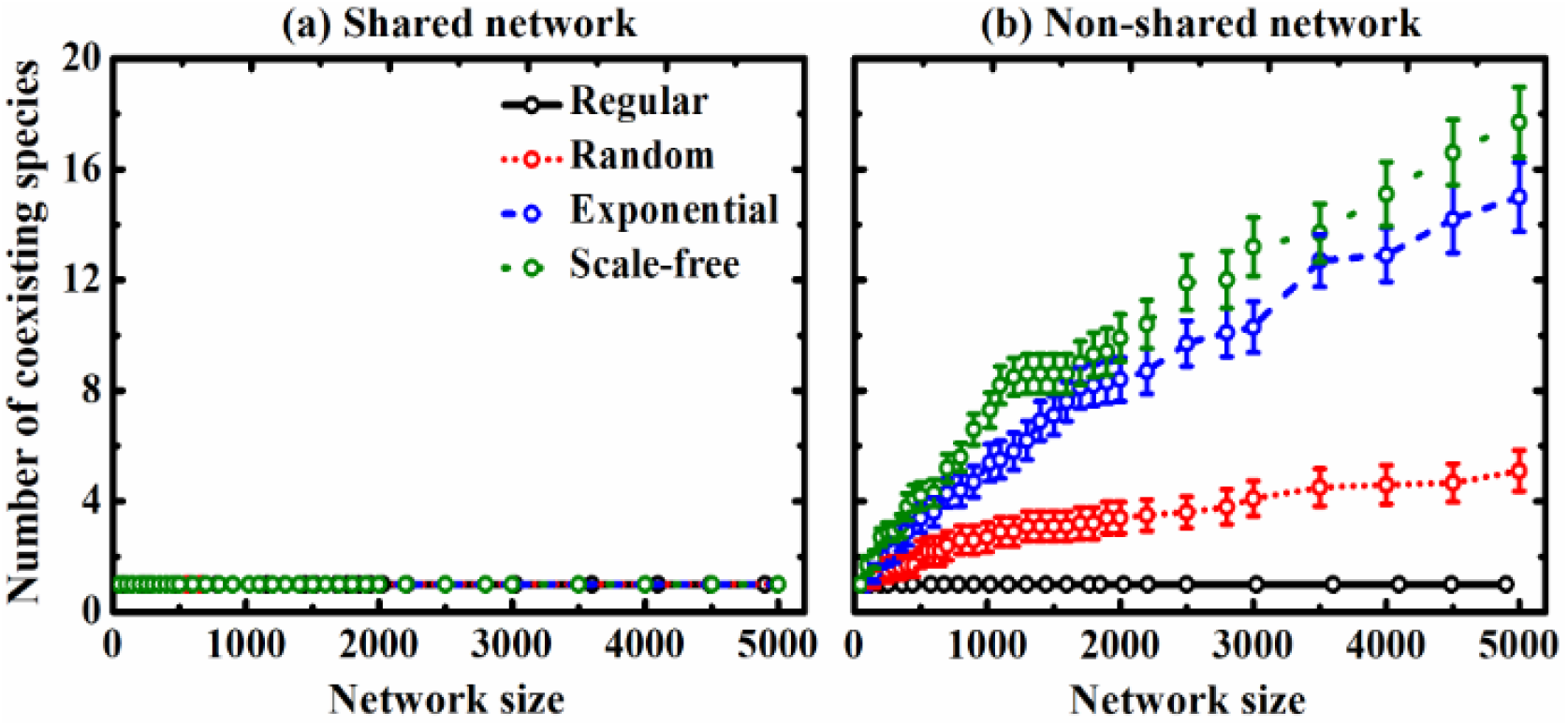
Species-area relationship between network size (i.e. total number of patches) and the number of coexisting species at steady state in hierarchical competitive metacommunities with preemptive competition (mean ± SD of 100 replicates). To exclude the colonization-competition tradeoff, all species are assumed to have the same colonization and extinction rate (*c*=*e*=0.05) in (a) shared vs. (b) non-shared networks (i.e. species-specific dispersal but with the same heterogeneity), with fixed average patch linking degree 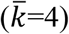. Again, four types of dispersal networks with contrasting heterogeneities are included: regular, random, exponential and scale-free networks.

## Results

### Two-species system

To get insights into the competitive dynamics, we first simply analyze two species (*A* – inferior competitor and *B* – superior competitor) competing for an empty patch *i* locally, thus the probability of the superior species *B* successfully colonizing the empty patch should be

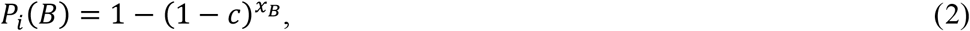

with 0< *c* <1. But if the species *B* fails to establish in there, then the inferior species *A* would have the opportunity to occupy this empty patch *i* with

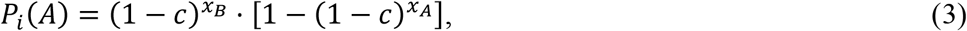

where *x_A_* (or *x_B_*) is the number of species *A* (or *B*) directly linked to the patch *i*.

Now we analyze if there exists the possibility that the inferior species *A* has higher opportunity to occupy the focal empty patch *i* than the superior species *B*. By setting *P_i_*(*A*) > *P_i_*(*B*), we have

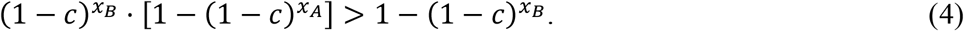

As such, the conditions for *P_i_*(*A*) > *P_i_*(*B*) can be derived as

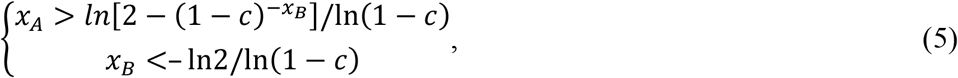

otherwise *P_i_*(*A*) > *P_i_*(*B*) (see phase diagram at Fig. S23 in *Appendix*). Thus, the inferior competitor can have local superiority to colonize the vacant patches relative to the superior competitor when the inferior species has more dispersal links to those patches, indirectly demonstrating that both inferior and superior species might coexist regionally in non-shared dispersal networks with heterogeneity.

We then numerically simulate the coexistence of two competitors with the same demography (i.e. identical colonization and extinction rate) in shared vs. non-shared dispersal networks with contrasting heterogeneities, including (from most homogeneous to most heterogeneous) regular, random, exponential and scale-free networks (Fig. 1). In general, both species cannot coexist regionally in the shared networks regardless of dispersal heterogeneity, as the superior species eventually excludes the inferior species (Fig. 1I-IV). In contrast, when their dispersal networks are non-shared (but with the same heterogeneity; e.g. scale-free networks in Fig. S3), they can co-occur stably (Fig. 1VI-VIII), except in the regular networks (analogous to “shared” homogeneous networks) where the poor competitor is excluded (Fig. 1V; see coexistence pattern in Fig. S4 in *Appendix*). Interestingly, increasing dispersal heterogeneity in the non-shared networks decreases the amplitude of stochastic fluctuations in patch dynamics and makes the species abundances converge (increased abundance of the inferior species and vice versa).

The coexistence patterns described above can, however, be altered by varying the species’ relative extinction or/and colonization rates (Fig. 2; Figs S5-S13 in *Appendix*) as well as the average linking degree (Fig. S14 in *Appendix*). On the one hand, both species expectedly display a monotonic decline in global patch occupancy (mean ± SD of 100 replicates) as the relative extinction rate (*e*/*c*) increases, thus speeding up species exclusion. This occurs irrespective of whether the dispersal networks are shared or not. Yet, species coexistence in the non-shared networks with higher heterogeneity can tolerate much higher *e*/*c*-ratios. On the other hand, increasing the relative colonization rate (*c/e;* i.e. dispersal rate) always leads to higher abundance of the superior species than the inferior species. In the shared networks, the inferior species is always outcompeted regardless of heterogeneity. In the non-shared networks but with the same heterogeneity, the inferior species remains longer in the system with increasing values of *c/e* when the network is heterogeneous. In this case, the patch occupancy of the inferior species initially increases and later declines to extinction, contrary to regular networks where the inferior species always goes extinct. Intermediate levels of *c/e* thus maximize the inferior species’ abundance and consequently promote species coexistence, as opposed to lower or higher dispersal rate which would speed up species exclusion. This outcome is similar to the case where the average linking degree is increased (Fig. S14 in *Appendix*). Furthermore, we find that increasing network heterogeneity increases the parameter space (*c/e*) for spatial coexistence.

We further explore species coexistence when both competitors (again with the same demography) use dispersal networks with different heterogeneities (Fig. 3). If the poor competitor has a regular dispersal network, the strong competitor with higher dispersal heterogeneity (including random, exponential and scale-free) can outcompete it (Fig. 3II-IV). In contrast, the inferior species with higher dispersal heterogeneity surprisingly can exclude the superior species with regular dispersal. In other words, dispersal heterogeneity can compensate for competitive disadvantage to some degree, and overturn the competitive outcome (Fig. 3I, V & IX). In other cases where both competitors exhibit dispersal networks with different heterogeneities (except regular networks), we observe that both species can co-occur stably, and higher dispersal heterogeneity decreases the fluctuation amplitude of patch occupancy dynamics at steady state (Fig. 3VI-VIII vs. X-XII).

### Multispecies system

Then we investigate how many competitors (with the same demography) can coexist stably in multispecies communities by varying the relative extinction and/or colonization rates, again considering shared vs. non-shared dispersal networks (Fig. 4; Figs S19-S20 in *Appendix*). Generally, increasing the relative extinction rate (*e/c*) reduces species richness in both shared and non-shared dispersal networks, but biodiversity maintenance differs depending on dispersal heterogeneity (Fig. 4a & c). When dispersal networks are shared, only the best competitor survives regardless of dispersal heterogeneity. In non-shared networks (but with the same heterogeneity), however, higher dispersal heterogeneity maintains more species (scale-free > exponential > random *>* regular). When varying the relative colonization rate (*c*/*e*), we find that again, only the best competitor can survive in shared networks regardless of dispersal heterogeneity, whereas more species can coexist with increasing dispersal heterogeneity in non-shared networks. Similar to increasing average linking degree (Fig. S21 in *Appendix*), the biodiversity response to increasing *c/e* is unimodal in the non-shared heterogeneous networks: moderate levels of *c/e* maximize species diversity, while lower or higher dispersal rates result in more species extinctions.

Finally we examine the effect of network size on biodiversity maintenance (i.e. the species-area curve), again in both shared and non-shared dispersal networks with contrasting heterogeneities (Fig. 5). In shared networks, only the best competitor survives regardless of network size and dispersal heterogeneity. In contrast, an increasing network size in non-shared heterogeneous networks (except in the regular network where only the best competitor survives) surprisingly leads to a monotonic increase in species richness, with greater species richness at higher dispersal heterogeneity.

## Discussion

Our spatially explicit model focuses on how species-specific dispersal networks and hierarchically preemptive competition interact in affecting species coexistence, assuming that all species have the same demographic traits to exclude the colonization-competition tradeoff. Yet, current theoretical understanding mostly comes from models with very regular connections among patches, in contrast with dispersal heterogeneity in natural systems which is far from regular (Hanski & Ovaskainen 2000; Fortuna et al. 2006; McIntire et al. 2007). In our study, we thus concentrate on dispersal heterogeneity, specifically in network structure (including regular, random, exponential and scale-free), to demonstrate the importance of dispersal heterogeneity for spatial coexistence. We indeed observe that, regardless of dispersal heterogeneity, species cannot coexist stably in shared networks if there is no colonization-competition tradeoff (Figs 1, 2 & 4). Sharing the same dispersal pathways implies that competitors encounter each other very frequently, so the best competitor would eventually drive all other species to extinction by quickly seizing the empty patches. Thus, previous patch-dynamic models focusing only on shared regular or random networks, might have largely underestimated species diversity, as species in natural communities always exhibit diverse dispersal patterns with more or less heterogeneity.

In contrast to shared dispersal networks, non-shared heterogeneous networks (even at low levels of heterogeneity) greatly promote spatial coexistence and therefore biodiversity maintenance, especially at higher dispersal heterogeneity (Figs 1, 2 & 4). We further explore the mechanism underlying these competitive outcomes by analyzing the spatial distribution for each species subject to its specific dispersal network (Fig. S4 in *Appendix*), and relating a patch’s incidence (i.e. the proportion of time steps that the node is occupied along the dynamics) with its linking degree as well as with the average linking degree of the patches it interacts with (Figs S15-S18 in *Appendix*). As observed in Fig. S4, species-specific dispersal networks with heterogeneity allow species to locally form many self-organized clusters of occupied patches, with the most connected patches as the core. This can be ascribed to variability in linking degree across patches, which results in variability across patches’ incidence (Figs S15-S16 in *Appendix*). Obviously, patch incidence grows non-linearly with linking degree (Eq. 1), that is, patches require a minimum linking degree to stay occupied in the majority of time steps, above which patch incidence saturates. In turn, the highly connected patches for a focal species can provide benefit for their directly linked patches in terms of incidence (Figs S17 & S18 in *Appendix*). For example, when comparing patches with the same linking degree, those attached to the more well-connected patches have a higher incidence (positive feedback). Specifically, conspecifics tend to develop into clusters segregated from other species in space because of dispersal heterogeneity, which increases the frequency of neighbourhood dispersal within conspecifics and decreases competition among heterospecifics, thereby allowing demography-equivalent species to co-occur regionally in spite of preemptive competition (so-called the segregation-aggregation mechanism; Pacala 1997; Murrell et al. 2001; Holyoak & Loreau 2006). This mechanism can also be thought of as generating a type of spatial refugia for the poor competitors, i.e. locations within the clusters favoring persistence of the focal species after the extinction in surrounding areas. Alternatively, spatially different dispersal network connectivities can result in local advantages for each species in terms of patch re-colonization, for example, the inferior species within a local area containing highly connected patches, can have higher effective colonization rate due to higher number of links for the core patches. Essentially, such fundamental mechanism that promotes regional coexistence belongs to spatial heterogeneity in population growth rate. Therefore, the non-shared heterogeneous dispersal networks offer a new way that the classic coexistence mechanism of spatial heterogeneity can occur.

When two competitors differ in dispersal heterogeneity, they are able to coexist stably even though the poor competitor displays lower dispersal heterogeneity than the strong competitor (Fig. 3), again confirming that dispersal heterogeneity can weaken interspecific competition and therefore promote species coexistence. Interestingly, this prediction was observed empirically by Yeaton & Bond (1991), where two competing shrub species with dispersal differences (one with ant-dispersed seeds and another with wind-dispersed seeds) can co-occur stably. Furthermore, a poor competitor with higher dispersal heterogeneity can even exclude a strong competitor with regular (homogeneous) dispersal, thus altering the species’ competitive rankings. Essentially, relative to the homogeneous dispersal of the superior competitor, even very low dispersal heterogeneity (e.g. random dispersal) can greatly promote local recolonization opportunities of the poor competitor, resulting in a large numerical advantage that overwhelms the competitive superiority and therefore excludes the strong competitor. This indicates that effects of spatial dispersal heterogeneity far outweigh the effect of preemptive competition, further confirming its important role in maintaining species diversity. However, these patterns of regional coexistence are greatly mediated by species life-history traits. Interestingly, increasing species dispersal (rates or links) prompts a unimodal biodiversity response in non-shared heterogeneous networks, demonstrating that effects of species dispersal on multispecies coexistence can be positive as well as negative (Figs 2 & 4 and Fig. S21 in *Appendix*). It is intuitive that a species is unable to persist locally at very low levels of dispersal rate. Yet, too much dispersal promotes patch colonization opportunities for the superior competitors, and consequently leads to the region-wide exclusion of the poor competitors, similar to increasing average patch linking degree (Figs S14 & S21 in *Appendix*).

A final observation is that increasing network size monotonically increases species richness in non-shared heterogeneous networks, with stronger species persistence at higher dispersal heterogeneity (Fig. 5). Essentially in the non-shared networks with heterogeneity, there is a much higher chance that species have exclusive access to specific patches whenever the landscape is composed of a very large number of patches. The resulting monotonically increasing species-area curves refute the previous view that the number of species coexisting cannot exceed the number of limited factors (Levin 1970; Tilman 1982). Instead, we theoretically demonstrate that, when there is species-specific dispersal heterogeneity, many more species than the number of limiting resources should be able to coexist, as empirically observed in several natural systems (Tilman 1982; Kotler & Brown 1988; Wellborn et al. 1996). Previously, coexistence of an unlimited number of species in a spatial context was ascribed to the colonization-competition tradeoff (Tilman 1994) rather than to spatial heterogeneity (Adler & Mosquera 2000). Our model provides an alternative explanation, if the landscape is large enough, for non-shared dispersal heterogeneities supporting the coexistence of many more species than expected, based on the segregation-aggregation mechanism.

By demonstrating that the architecture of dispersal networks strongly governs species coexistence, mediated by species life-history attributes, our work helps fill the gap between network structure and spatial competition. We find that incorporating species-specific dispersal heterogeneities into the traditional hierarchical competitive systems can greatly promote regional coexistence owing to the formation of self-organized clusters. This implies that traditional shared lattice- or randomly-structured models might have severely underestimated biodiversity maintenance. More importantly, the model suggests significant implications for biodiversity conservation and management. For instance, as different species often display diverse patterns of patch connectivity based on their dispersal traits, we should first construct and analyze dispersal networks independently for multiple target species, and then overlay or intersect the multiple networks to find locations that are important for these species, so as to design multispecies conservation planning (e.g. Bunn et al. 2000; Urban & Keitt 2001; Nicholson & Possingham 2006). However, two caveats should be addressed when applying our model to terrestrial ecosystems. Firstly, although there have been a large number of studies on scale-free graphs (Barabasi & Albert 1999), actual patch mosaics seem to not quite fit the definitions of such well-studied networks so that they tend to not include the extremely connected patches that characterize scale-free networks (Urban et al. 2009). Secondly, it may be inappropriate to apply a graph representation for some landscapes if habitat patches are poorly resolved spatially (Urban & Keitt 2001). For example, habitat quality varies continuously and subtly over the landscape, thus aggregating this variability into discrete patches would be inappropriate (e.g. Liao et al. 2013b). Overall, by integrating both network and metapopulation approaches, our modelling study provides a new way to understand the coexistence mechanism of spatial dispersal heterogeneity, thereby strengthening our comprehension of biodiversity maintenance in hierarchical competitive communities.

## Acknowledgements

This study was supported by the National Science Foundation of China (No. 31760172 & 31901175), the Thousand Young Talents Plan of China, the Key Joint Youth Project of Jiangxi Province (No. 20192ACBL21029), the Jiangxi Provincial Education Department (No. GJJ160274), and the Doctoral Scientific Research Foundation of Jiangxi Normal University (No. 12017778).

## Author contributions

J.L. conceived and wrote this manuscript, and H.Z. conducted simulations and analyzed results.

## Supplementary Material

Appendix accompanying this manuscript is supplied.

## Data accessibility

This is a theoretical modelling study which does not use data.

