## Appendix for "Non-shared dispersal networks with heterogeneity promote species coexistence in hierarchical competitive communities"

### Appendix – *Figures*

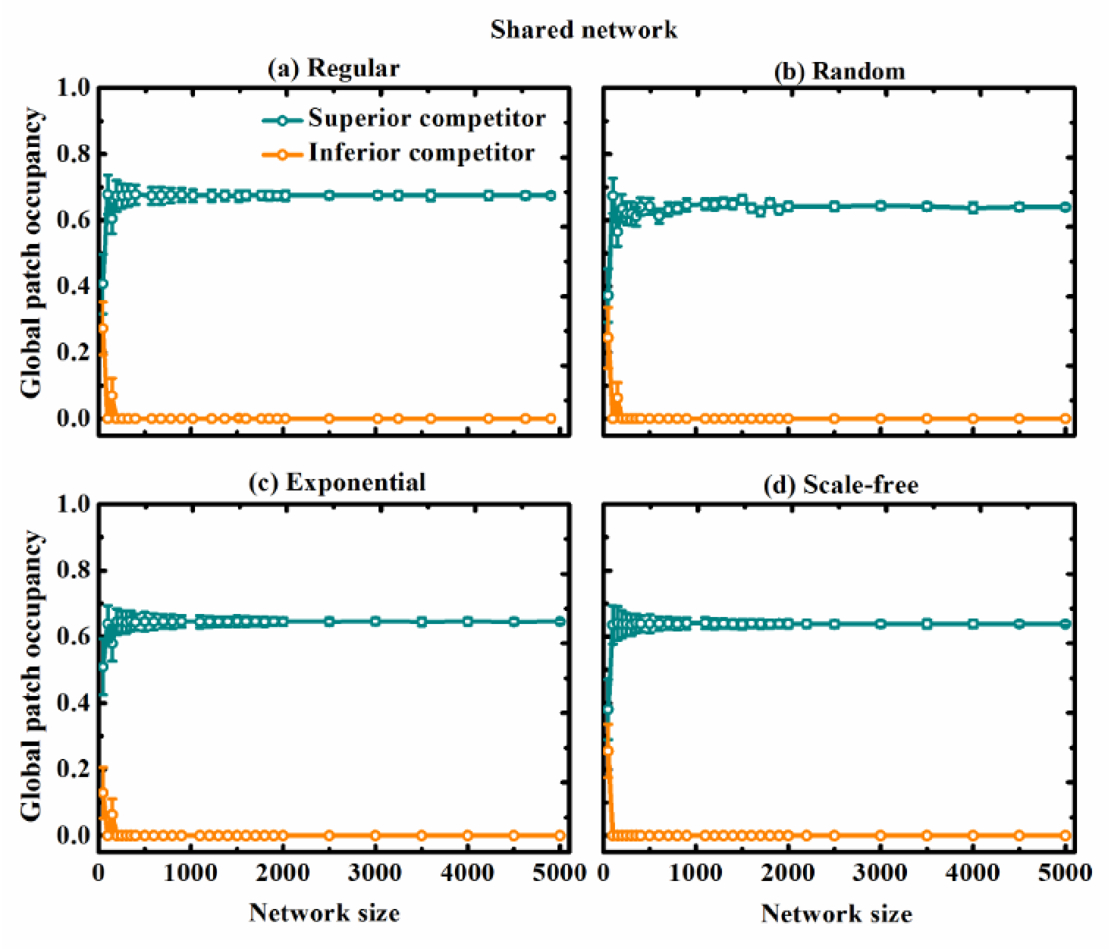


**Figure S1.** Effect of networks size on global patch occupancy (mean ± SD of 100 replicates) of two competing species at steady state in shared dispersal networks with the same average linking degree $\bar{k}$=4, including (a) regular, (b) random, (c) exponential and (d) scale-free networks. Parameter values for both species are the same: $c$=$e$=0.05.

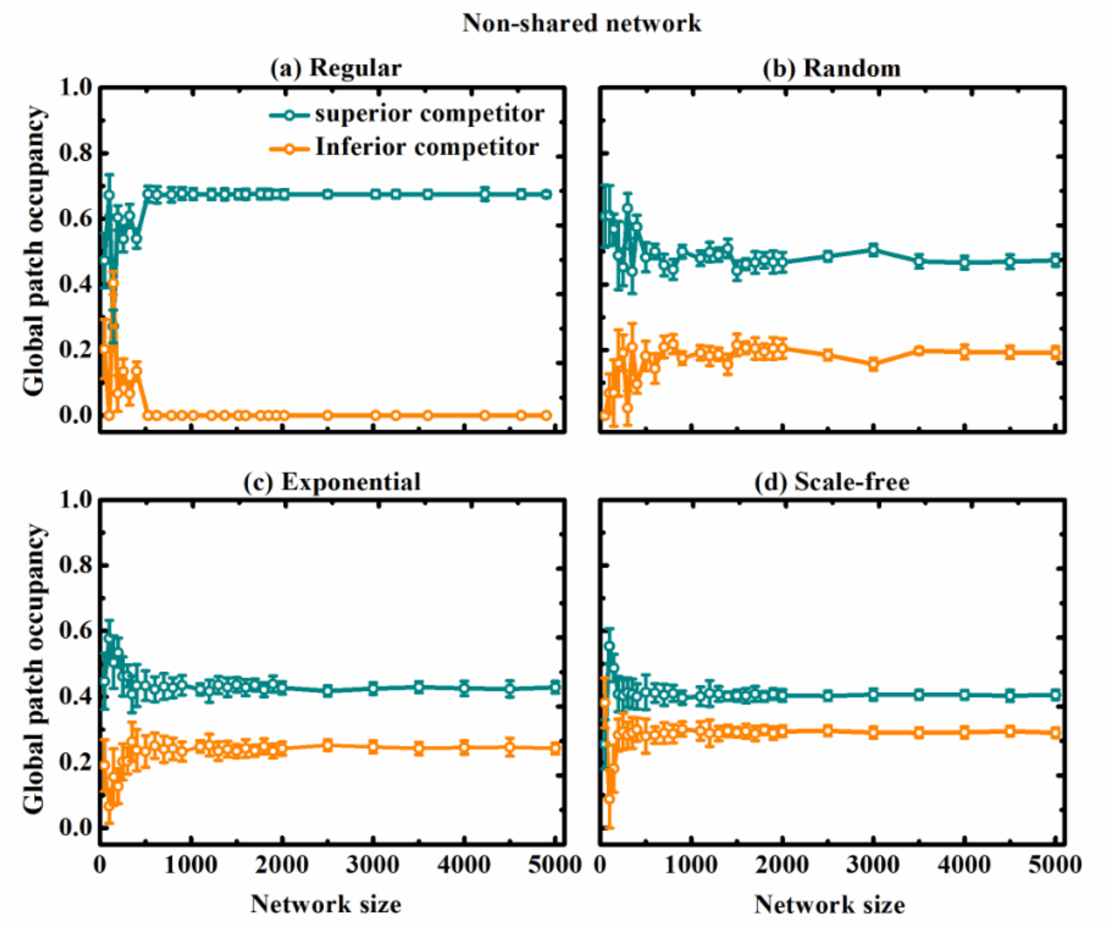


**Figure S2.** Effect of networks size on global patch occupancy (mean ± SD of 100 replicates) of two competing species at steady state in non-shared dispersal networks with the same average linking degree $\bar{k}$=4, including (a) regular, (b) random, (c) exponential and (d) scale-free networks. Parameter values for both species are the same: $c$=$e$=0.05.

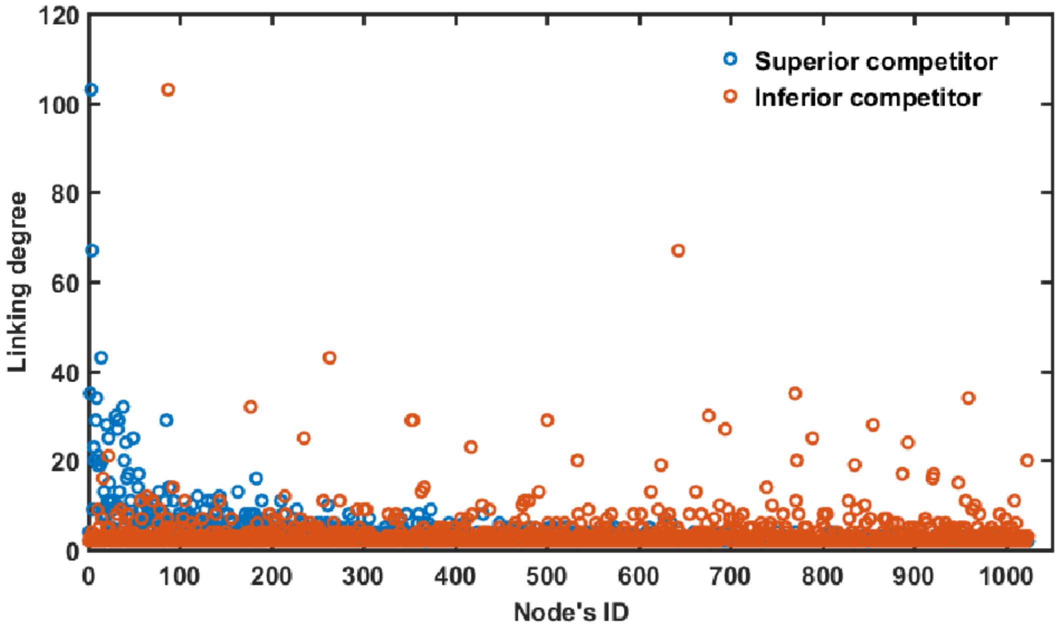


**Figure S3.** Patch linking degree distribution of two scale-free dispersal networks that are non-shared for two competitors, containing 1024 patches with 2048 links.

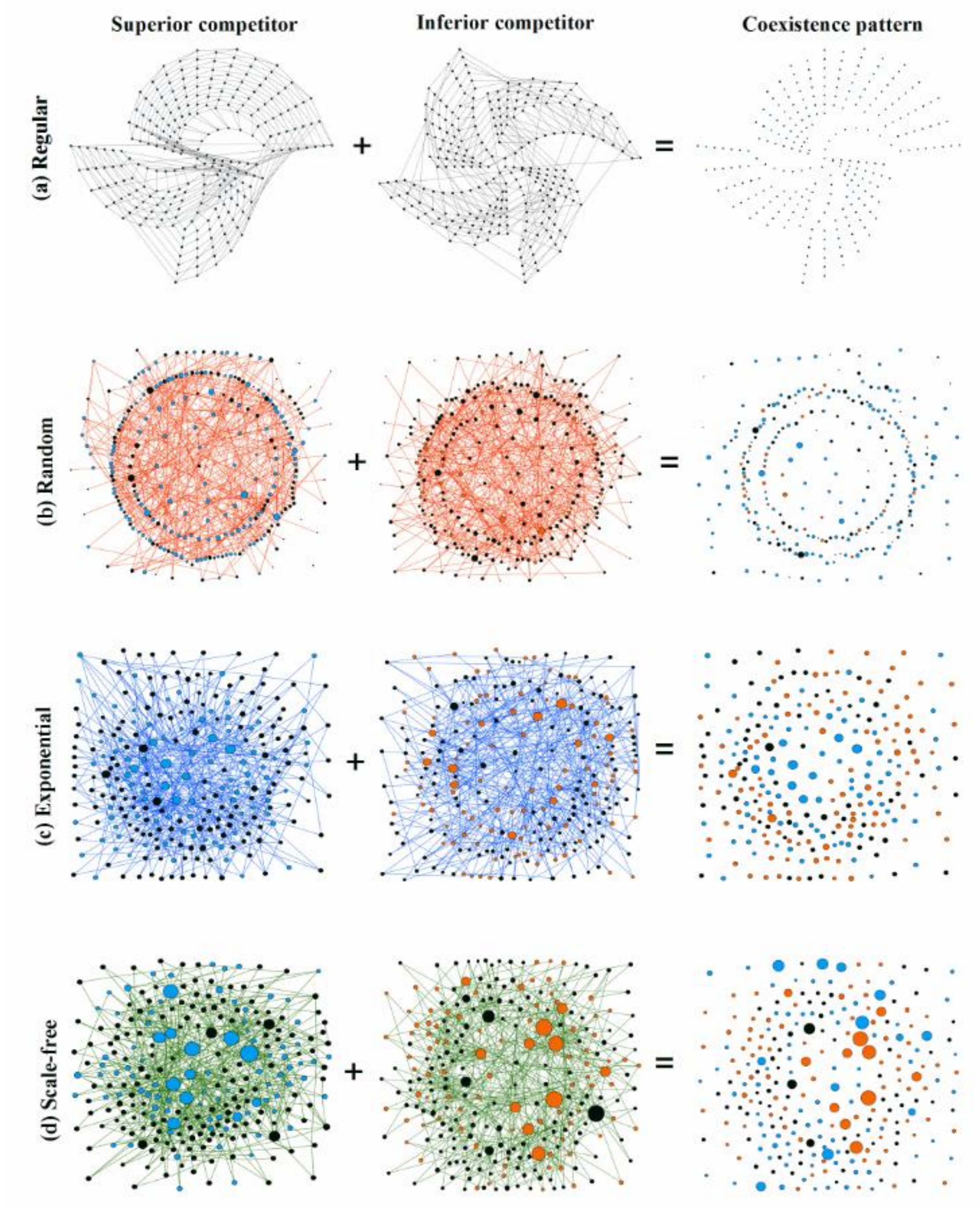


**Figure S4.** Coexistence pattern of two competing species at steady state ($t$=10,000) in non-shared dispersal networks (but with the same heterogeneity) of 256 patches with 512 links at fixed average patch linking degree $\bar{k} = 4$, including (a) regular, (b) random, (c) exponential and (d) scale-free networks. Patch size is proportional to its linking degree. Parameter values for both species are the same: $c$=$e$=0.05.

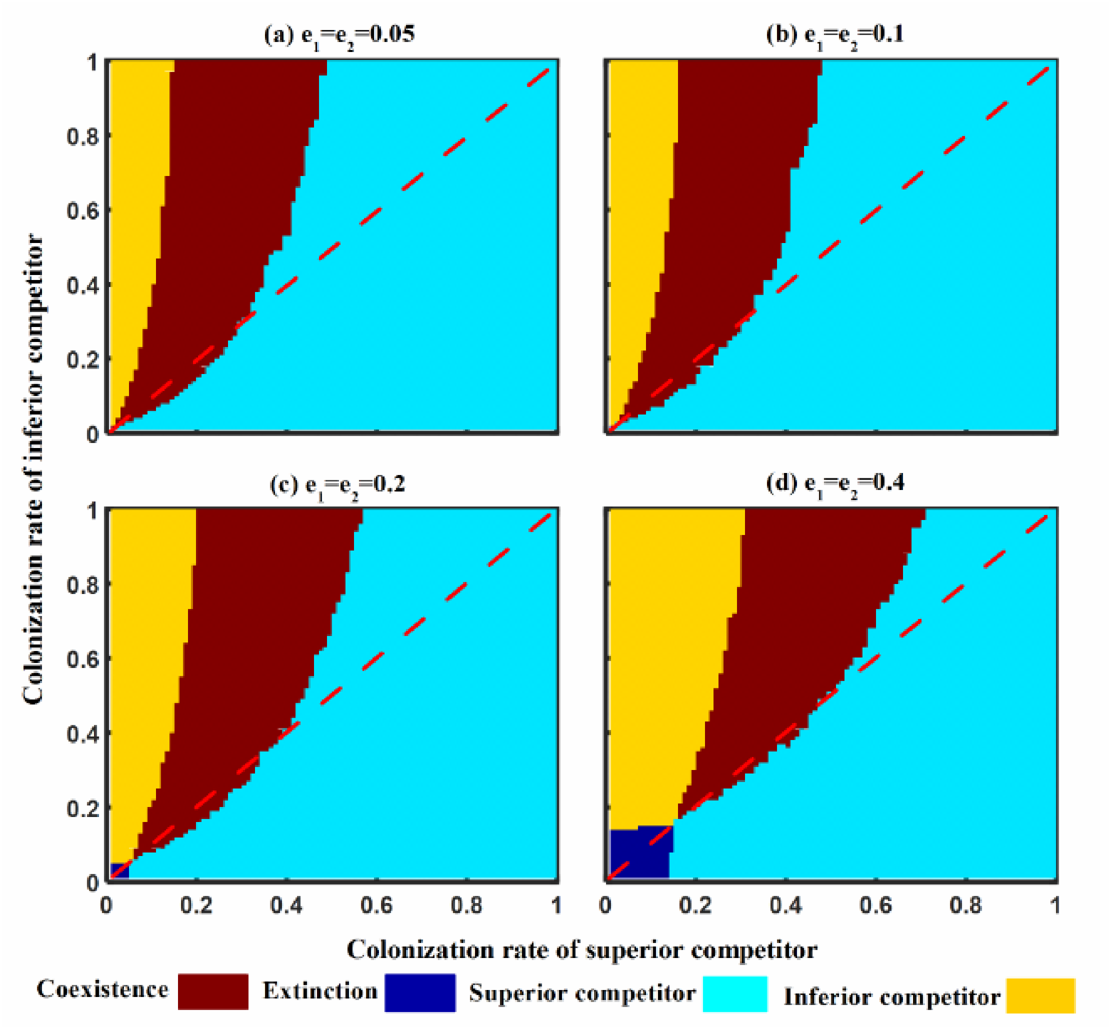


**Figure S5.** Interactive effects of variation in both species colonization rates on coexistence pattern at steady state in non-shared scale-free dispersal networks consisting of 1024 patches and 2048 links, simultaneously varying extinction rate for both species: (a) $e_1= e_2=0.05$, (b) $e_1= e_2=0.1$, (c) $e_1= e_2=0.2$ and (d) $e_1= e_2=0.4$.

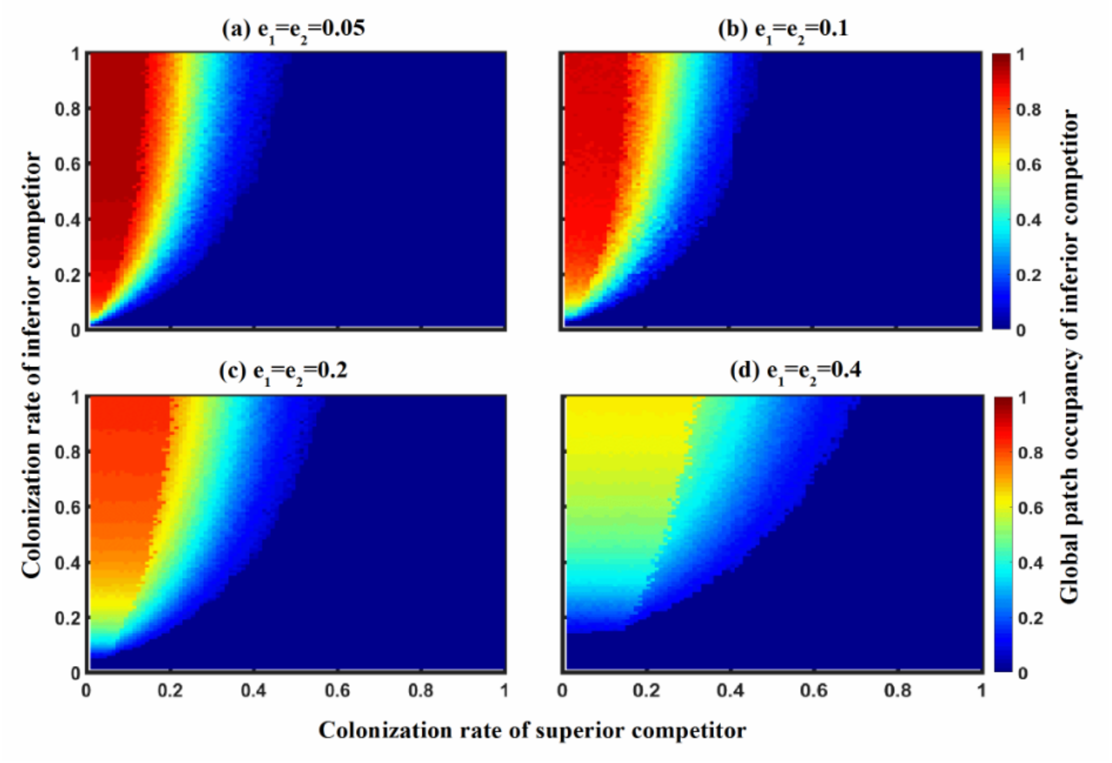


**Figure S6.** Interactive effects of variation in both species colonization rates on global patch occupancy of inferior competitor at steady state in non-shared scale-free dispersal networks consisting of 1024 patches and 2048 links, by varying species extinction rate for both competitors: (a) $e_1 = e_2 = 0.05$, (b) $e_1 = e_2 = 0.1$, (c) $e_1 = e_2 = 0.2$ and (d) $e_1 = e_2 = 0.4$.

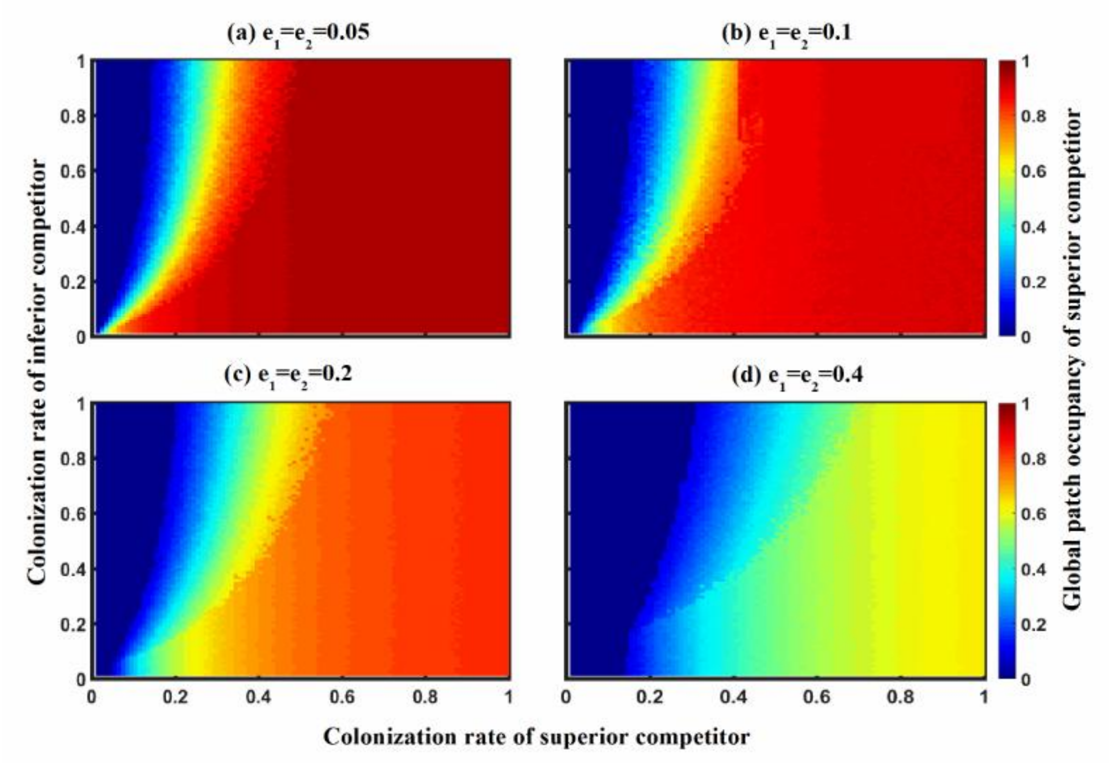


**Figure S7.** Interactive effects of variation in both species colonization rates on global patch occupancy of superior competitor at steady state in non-shared scale-free dispersal networks consisting of 1024 patches and 2048 links, by varying species extinction rate: (a) $e_1 = e_2$=0.05, (b) $e_1 = e_2$=0.1, (c) $e_1 = e_2$=0.2 and (d) $e_1 = e_2$=0.4.

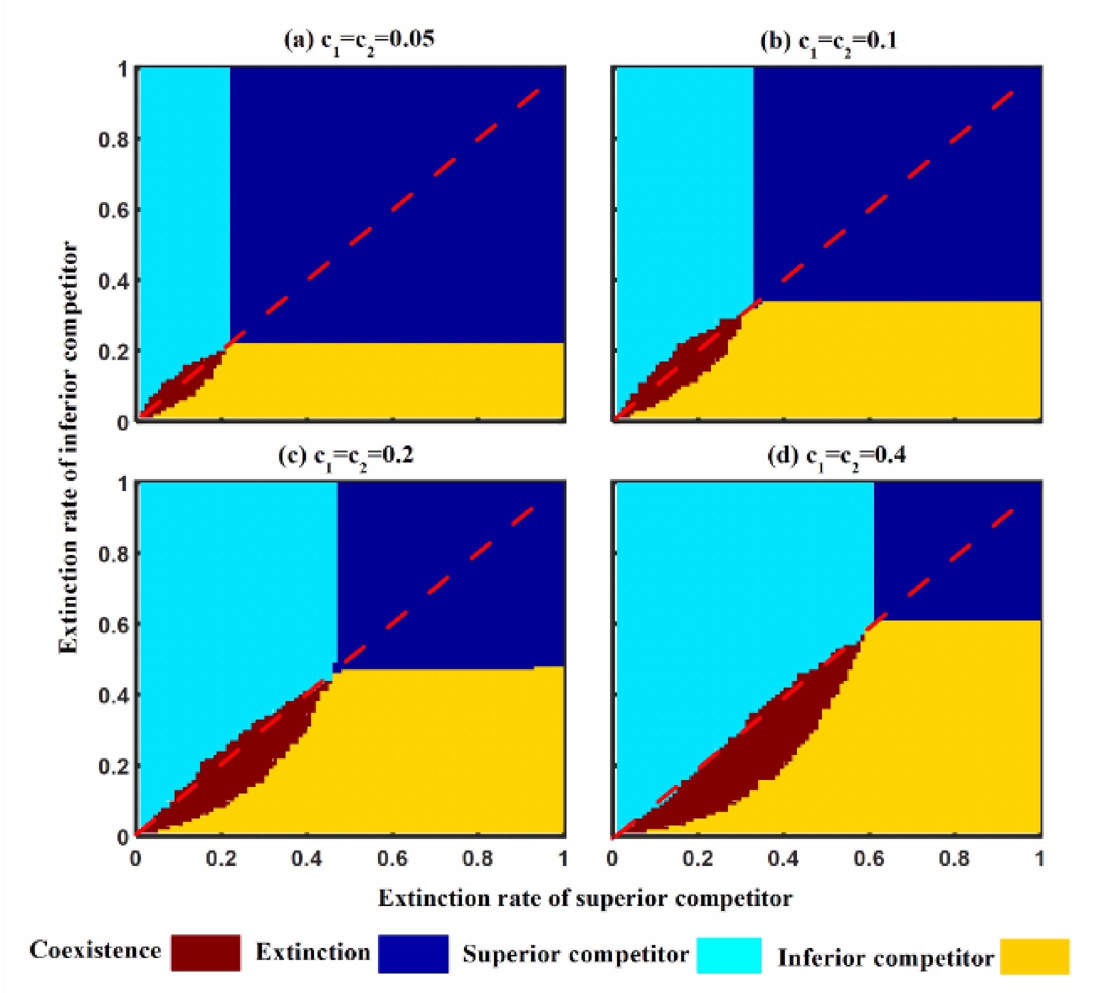


**Figure S8.** Interactive effects of variation in both species extinction rates on coexistence pattern at steady state in non-shared scale-free dispersal networks consisting of 1024 patches and 2048 links, simultaneously varying species colonization rate: (a) $c_1 = c_2 = 0.05$, (b) $c_1 = c_2 = 0.1$, (c) $c_1 = c_2 = 0.2$ and (d) $c_1 = c_2 = 0.4$.

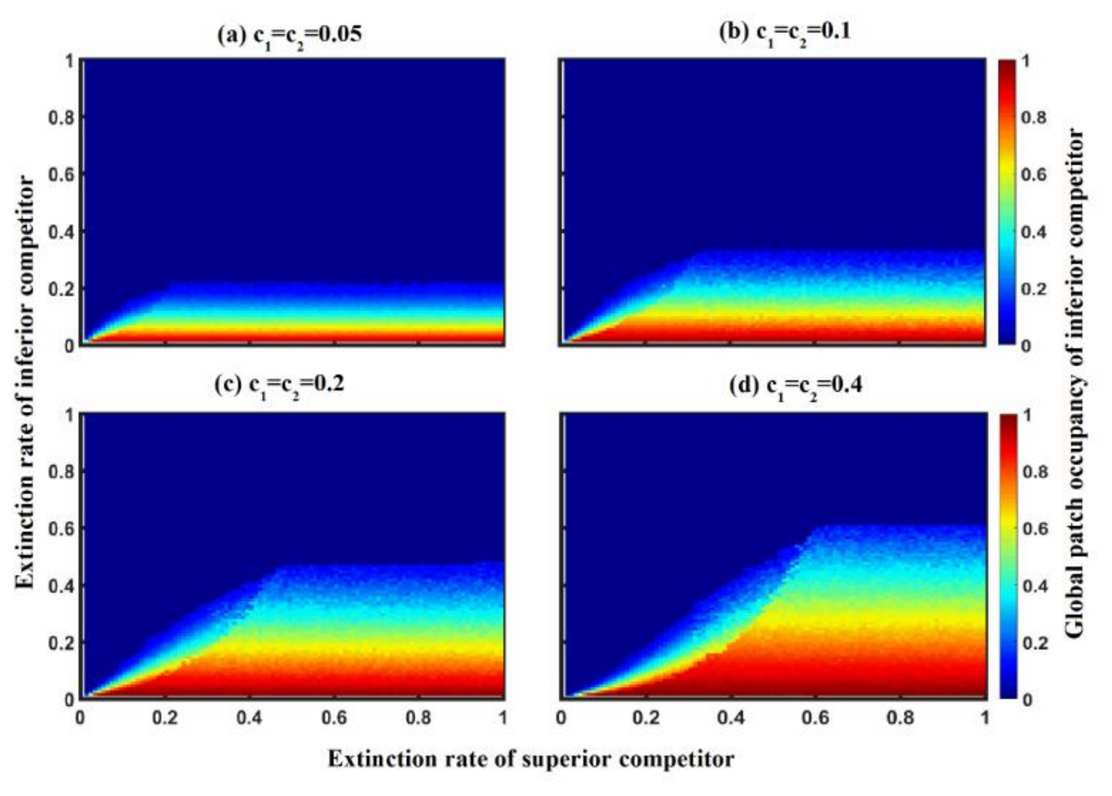


**Figure S9.** Interactive effects of variation in both species extinction rates on global patch occupancy of inferior species at steady state in non-shared scale-free dispersal networks consisting of 1024 patches and 2048 links, simultaneously varying species colonization rate: (a) $c_1 = c_2 = 0.05$, (b) $c_1 = c_2 = 0.1$, (c) $c_1 = c_2 = 0.2$ and (d) $c_1 = c_2 = 0.4$.

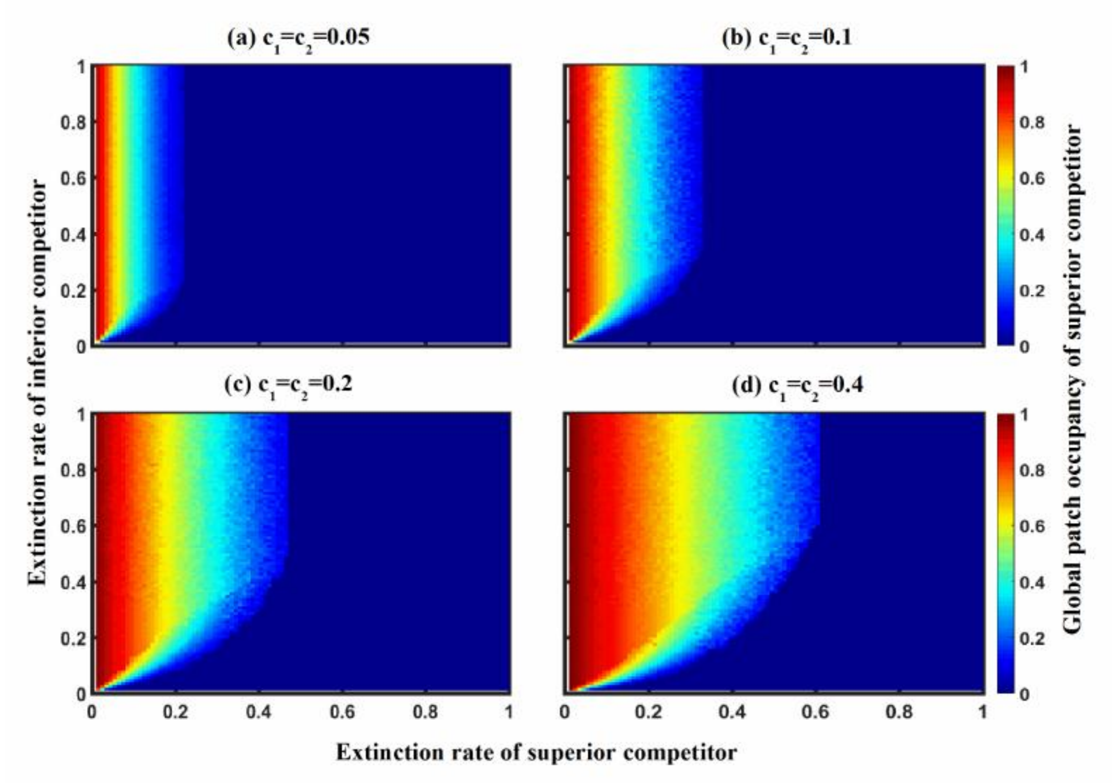


**Figure S10.** Interactive effects of variation in both species extinction rates on global patch occupancy of superior species at steady state in non-shared scale-free dispersal networks consisting of 1024 patches and 2048 links, simultaneously varying species colonization rate: (a) $c_1$= $c_2$=0.05, (b) $c_1$= $c_2$=0.1, (c) $c_1$= $c_2$=0.2 and (d) $c_1$= $c_2$=0.4.

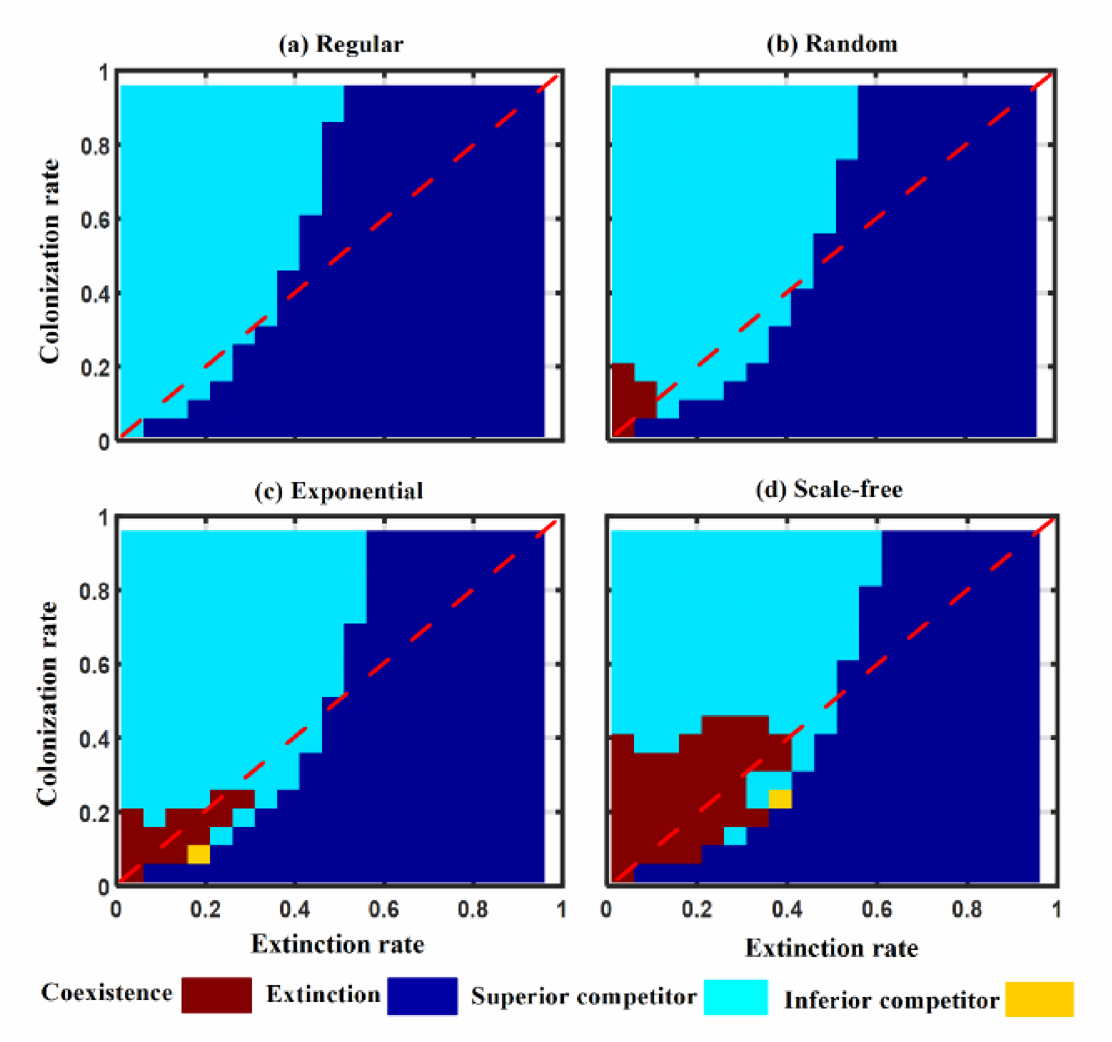


**Figure S11.** Interactive effects of varying both extinction and colonization rates for two competing species (with the same demographic traits) on coexistence pattern at steady state in the non-shared dispersal networks consisting of 1024 patches and 2048 links, including (a) regular, (b) random, (c) exponential and (d) scale-free networks.

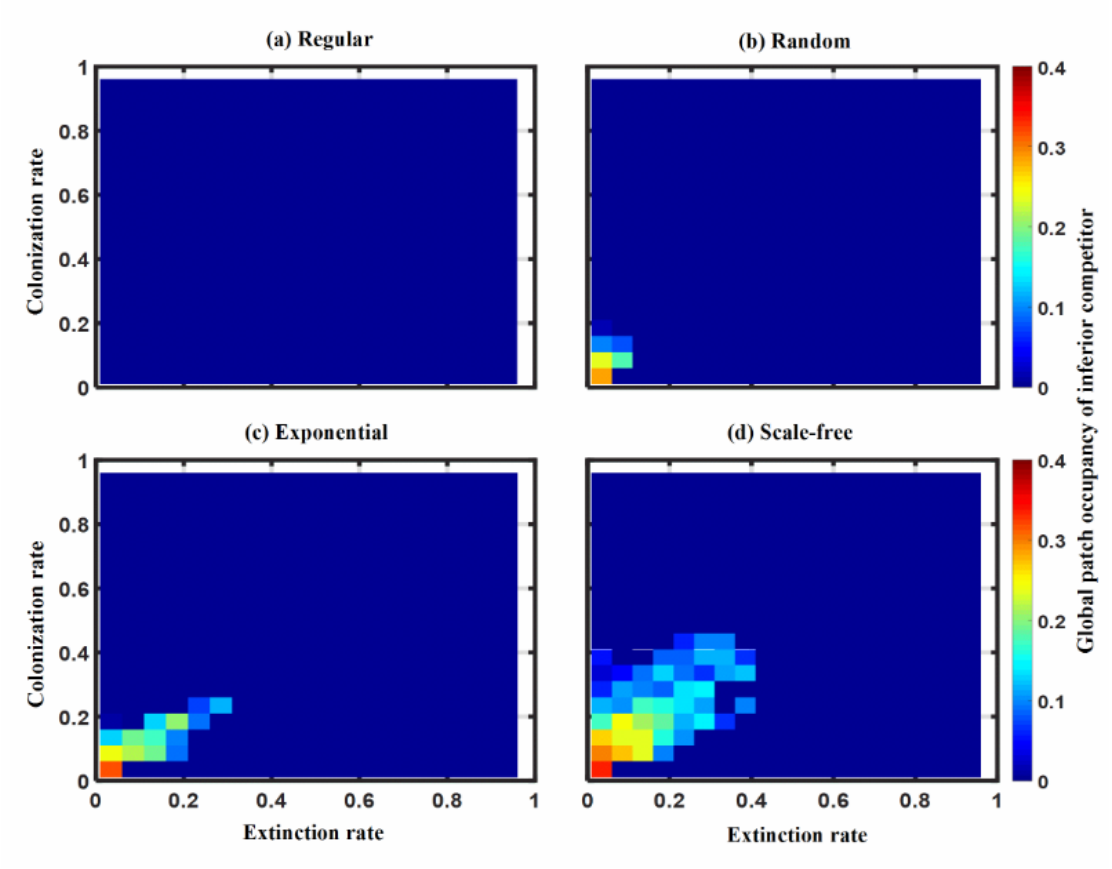


**Figure S12.** Interactive effects of varying both extinction and colonization rates for two competing species (with the same demographic traits) on global patch occupancy of inferior competitor at steady state in non-shared dispersal networks consisting of 1024 patches and 2048 links, including (a) regular, (b) random, (c) exponential and (d) scale-free networks.

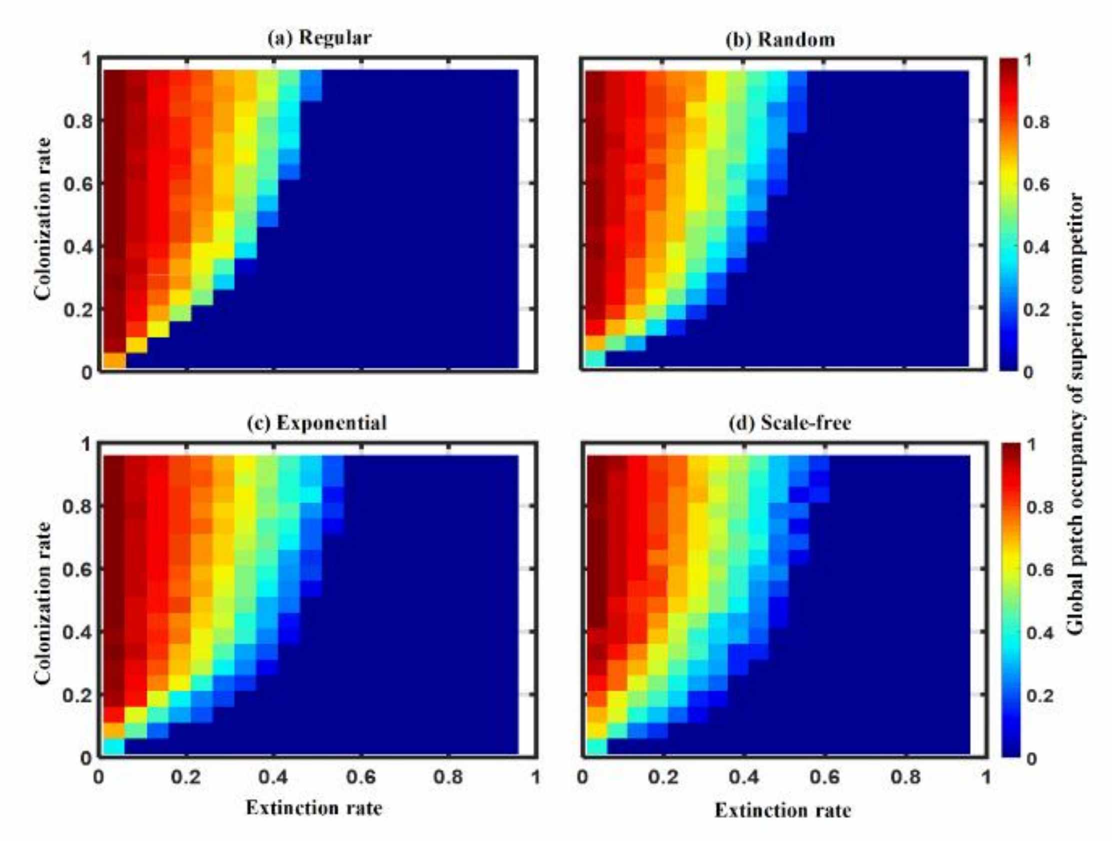


**Figure S13.** Interactive effects of varying both extinction and colonization rates for two competing species (with the same demographic traits) on global patch occupancy of superior competitor at steady state in non-shared dispersal networks consisting of 1024 patches and 2048 links, including (a) regular, (b) random, (c) exponential and (d) scale-free networks.

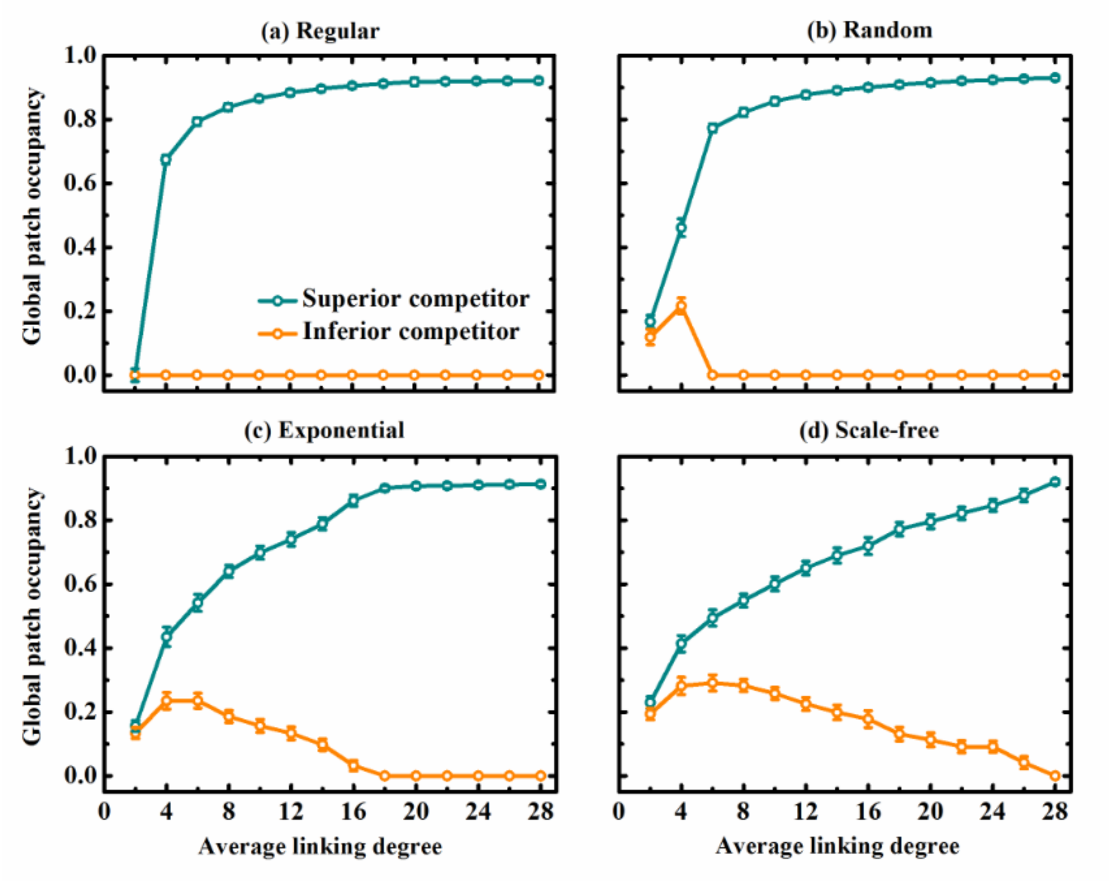


**Figure S14.** Effect of increasing average linking degree on the survival of two competing species (mean ± SD of 100 replicates) at steady state in the non-shared dispersal networks containing 1024 patches, including (a) regular, (b) random, (c) exponential and (d) scale-free networks. All species have the same demographic traits: $c$=$e$=0.05.

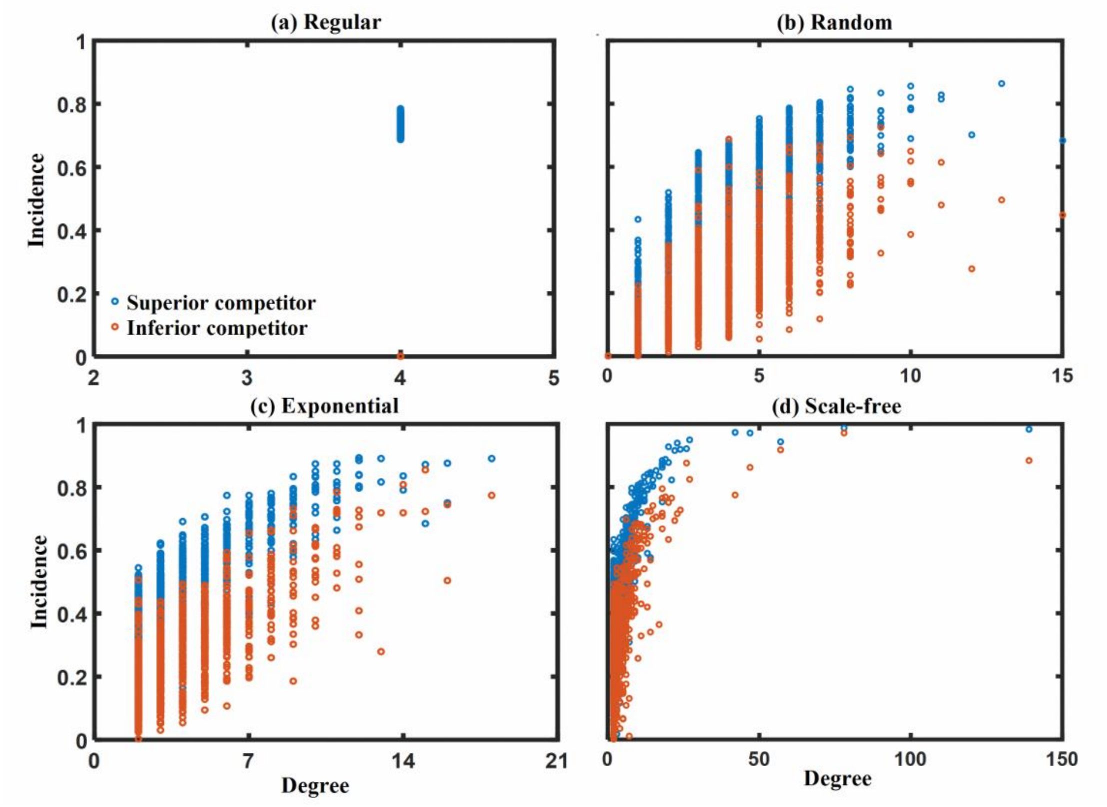


**Figure S15.** Relationship between patch incidence and its linking degree for two competitors in non-shared dispersal networks (but with the same heterogeneity) at steady state, containing 1024 patches and 2048 links. Four typical networks with contrasting heterogeneities are included: (a) regular, (b) random, (c) exponential and (d) scale-free networks. Incidence is calculated as the proportion of time steps (from 5000 to 10000 time steps) that a patch is occupied along the dynamics. Parameter values for both species are the same: $c=e=0.05$.

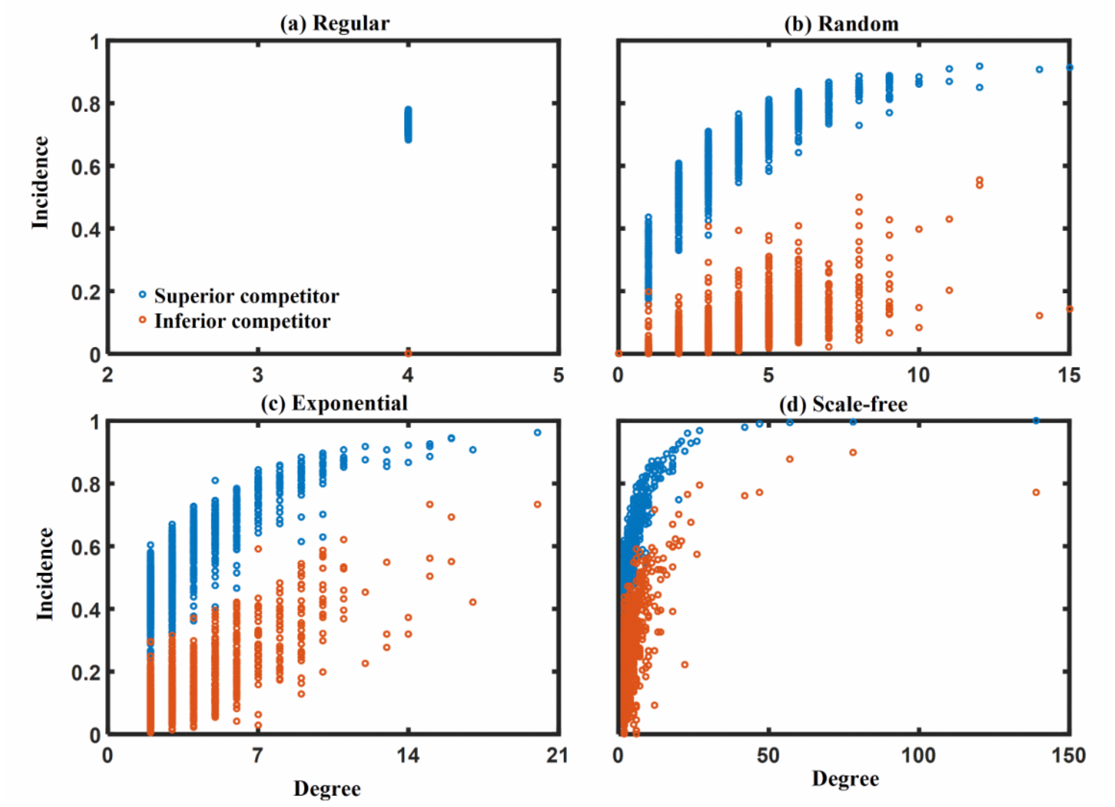


**Figure S16.** Relationship between patch incidence and its linking degree for two competitors in non-shared dispersal networks (but with the same heterogeneity) at steady state, containing 1024 patches and 2048 links. Four typical heterogeneous networks are included: (a) regular, (b) random, (c) exponential and (d) scale-free networks. Incidence is calculated as the proportion of time steps (from 5000 to 10000 time steps) that a patch is occupied along the dynamics. Parameter values for both species are the same: $c=e=0.1$.

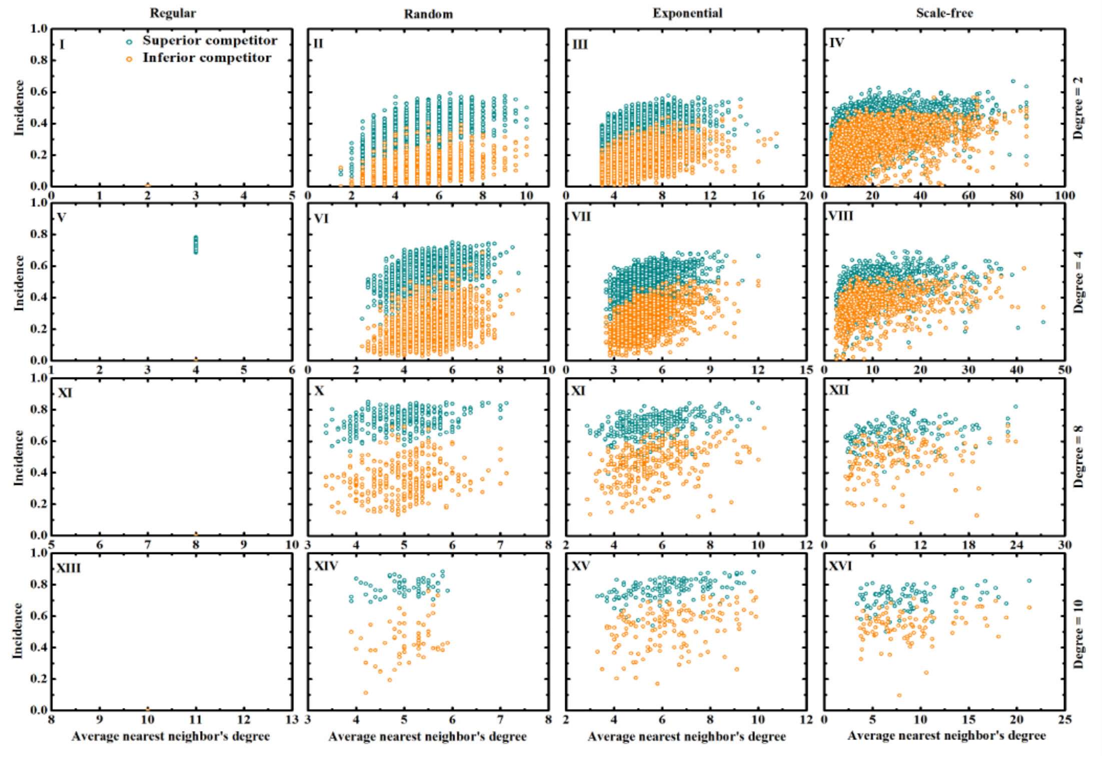


**Figure S17.** Patch incidence as a function of its average nearest neighbors' degree in two-competing system with non-shared dispersal networks (1024 patches and 2048 links) at steady state, for patches with four different linking degrees (*k*=2, 4, 8 & 10). Four typical dispersal networks with contrasting heterogeneities are considered: regular, random, exponential and scale-free networks. Topologically, the nearest neighbours of a focal patch are those patches directly linked to the latter. Incidence is calculated as the proportion of time steps (from 5000 to 10000 time steps) that a patch is occupied along the dynamics. Displayed results are obtained from the average dynamics of 10 replicates, with *c*=*e*=0.05.

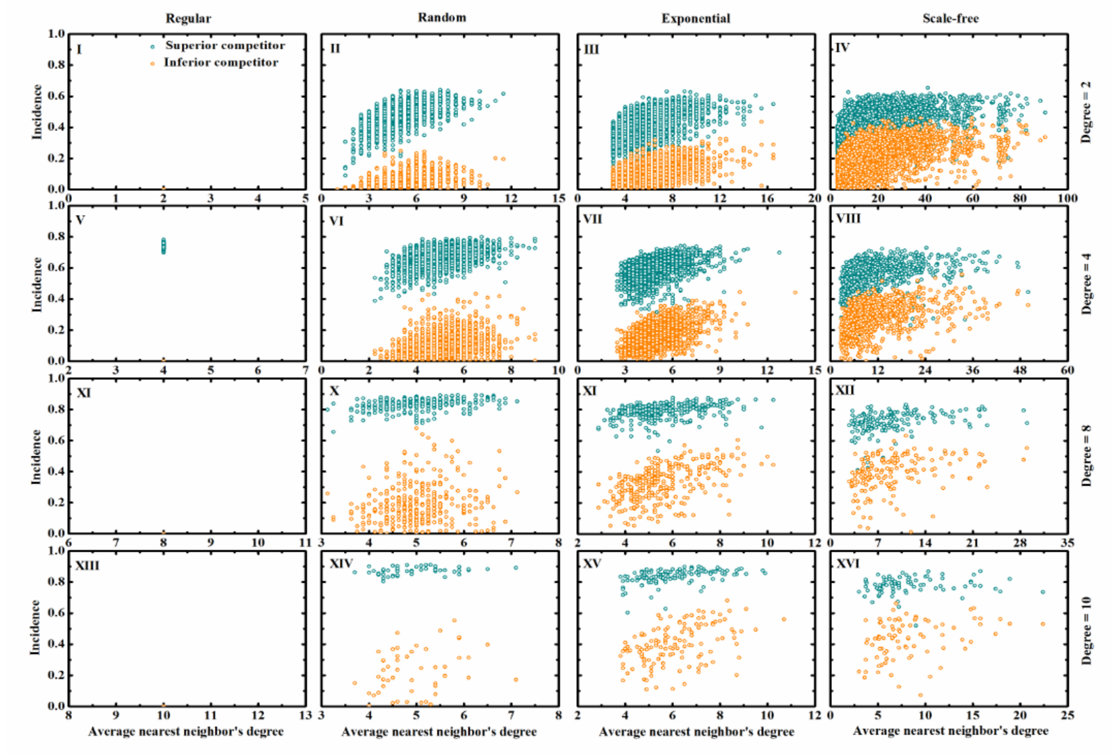


**Figure S18.** Patch incidence as a function of its average nearest neighbour's degree in two-competing system with non-shared dispersal networks (1024 patches and 2048 links) at steady state, for patches with four different linking degrees (*k*=2, 4, 8 & 10). Four typical dispersal networks with contrasting heterogeneities are considered: regular, random, exponential and scale-free networks. Topologically, the nearest neighbours of a focal patch are those patches directly linked to the latter. Incidence is calculated as the proportion of time steps (from 5000 to 10000 time steps) that a patch is occupied along the dynamics. Displayed results are obtained from the average dynamics of 10 replicates, with $c$=$e$=0.1.

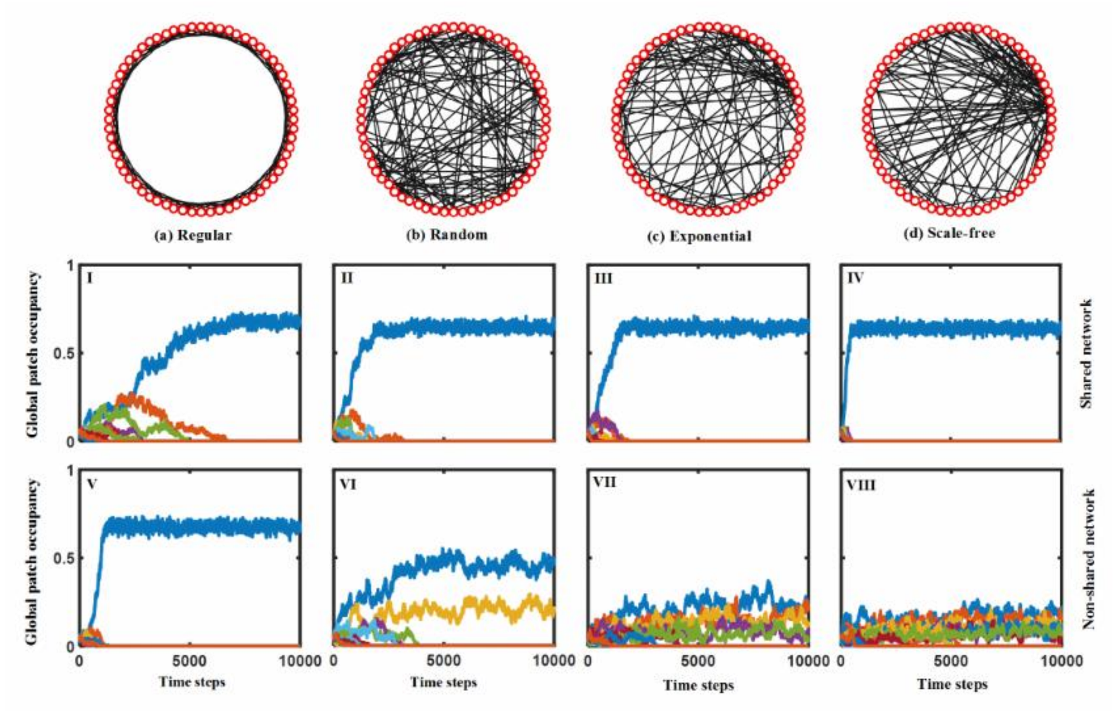


**Figure S19.** Patch dynamics of a hierarchical competitive metacommunity (initially consisting of 16 species) in shared vs. non-shared dispersal networks (containing 1024 patches and 2048 links) with contrasting heterogeneities: regular, random, exponential and scale-free networks. For display purposes, these networks in graphs (a-d) only consist of 64 patches and 128 links. Demographic trait values for all species are the same: $c$=$e$=0.05.

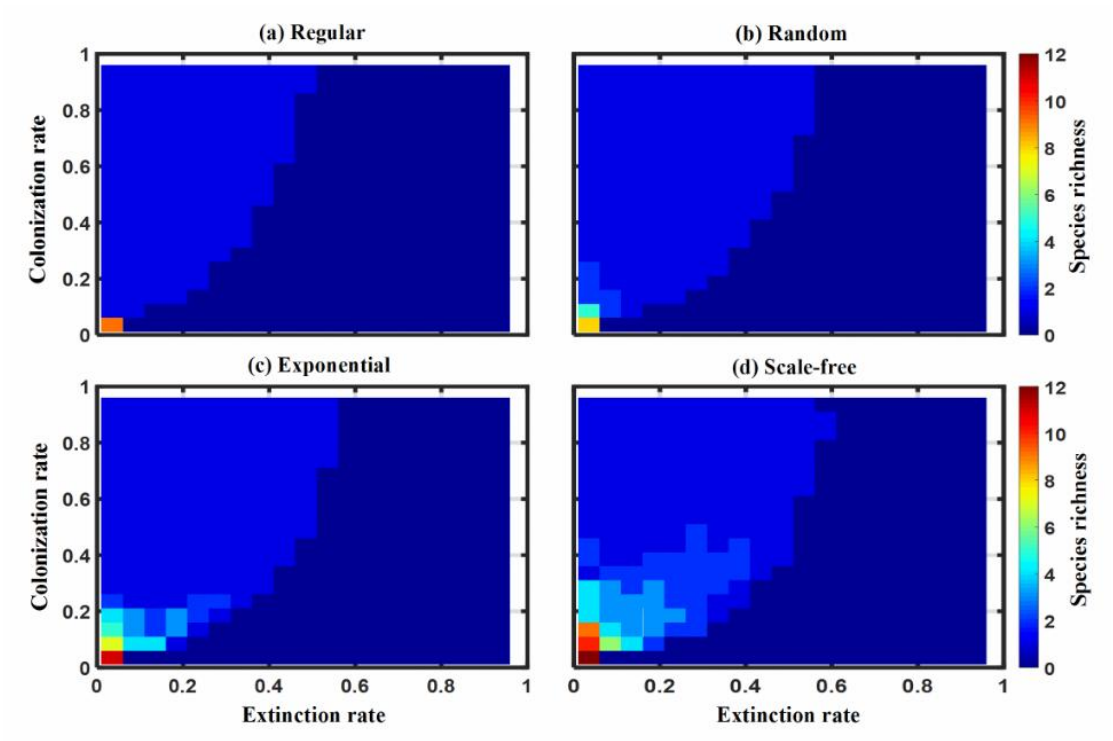


**Figure S20.** Interactive effects of species extinction and colonization rates on the number of coexisting species of hierarchical competitive metacommunities (initially containing 20 species) at steady state in non-shared dispersal networks containing 1024 patches and 2048 links, including (a) regular, (b) random, (c) exponential and (d) scale-free networks. All species have the same demographic traits: $c$=$e$=0.05.

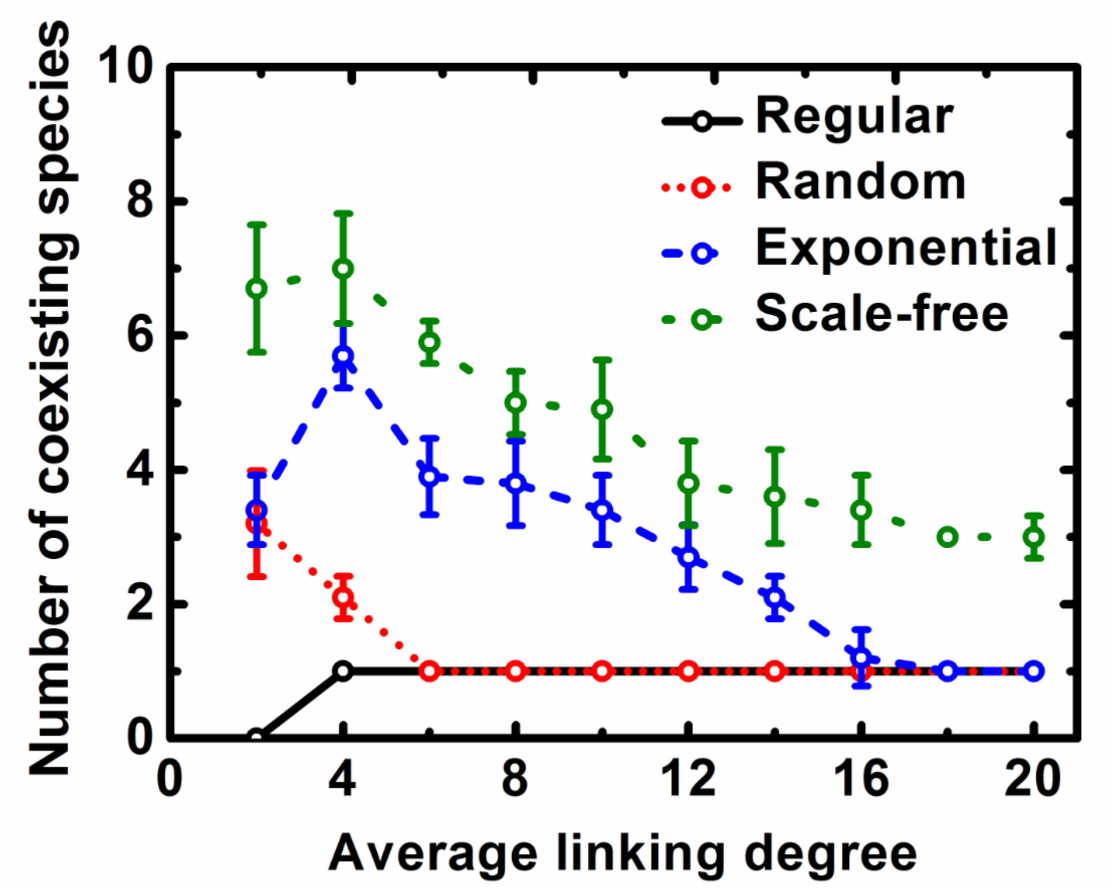


**Figure S21.** Effect of increasing average linking degree on the number of coexisting species (mean ± SD of 100 replicates) of hierarchical competitive metacommunities (initially containing 16 species) at steady state in non-shared dispersal networks containing 1024 patches, including regular, random, exponential and scale-free networks. All species have the same demographic traits: $c$=$e$=0.05.

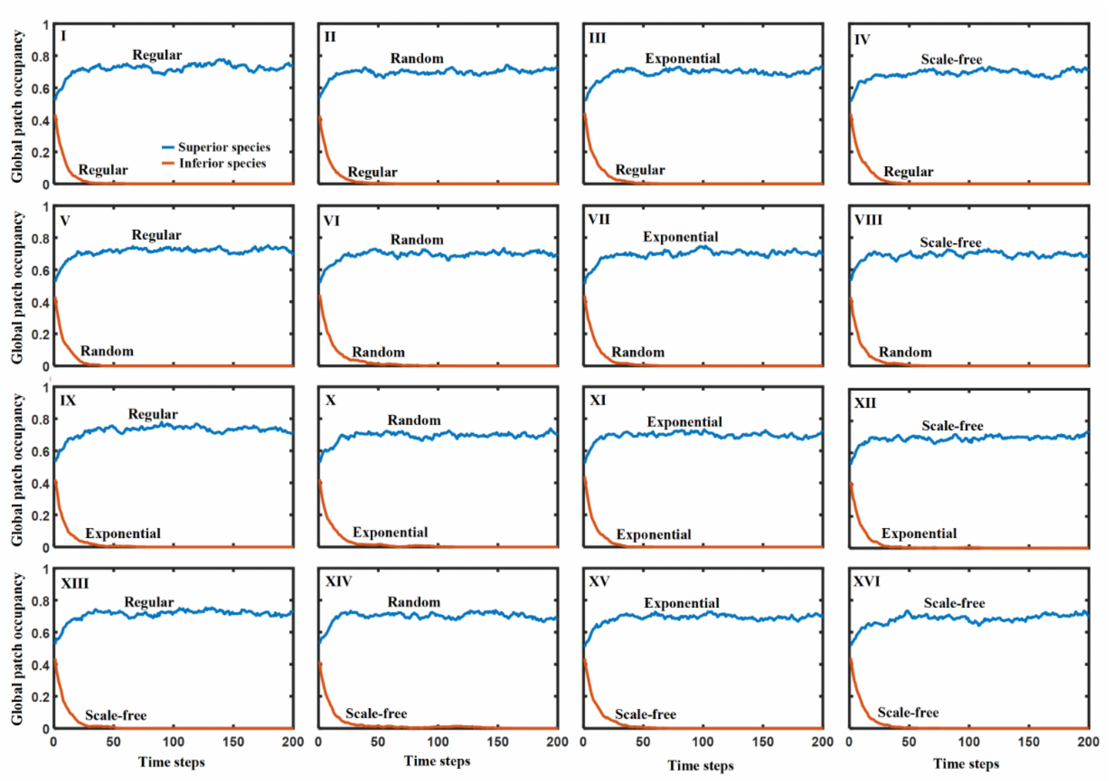


**Figure S22.** Patch dynamics of both inferior and superior competitors with different heterogeneous networks, consisting of 1024 patches and 2048 links. Here we consider the competitive displacement that the superior species can invade into and displace the inferior species. Four types of dispersal networks are included: regular, random, exponential and scale-free networks. Parameter values for both species are the same: $c=e=0.05$.

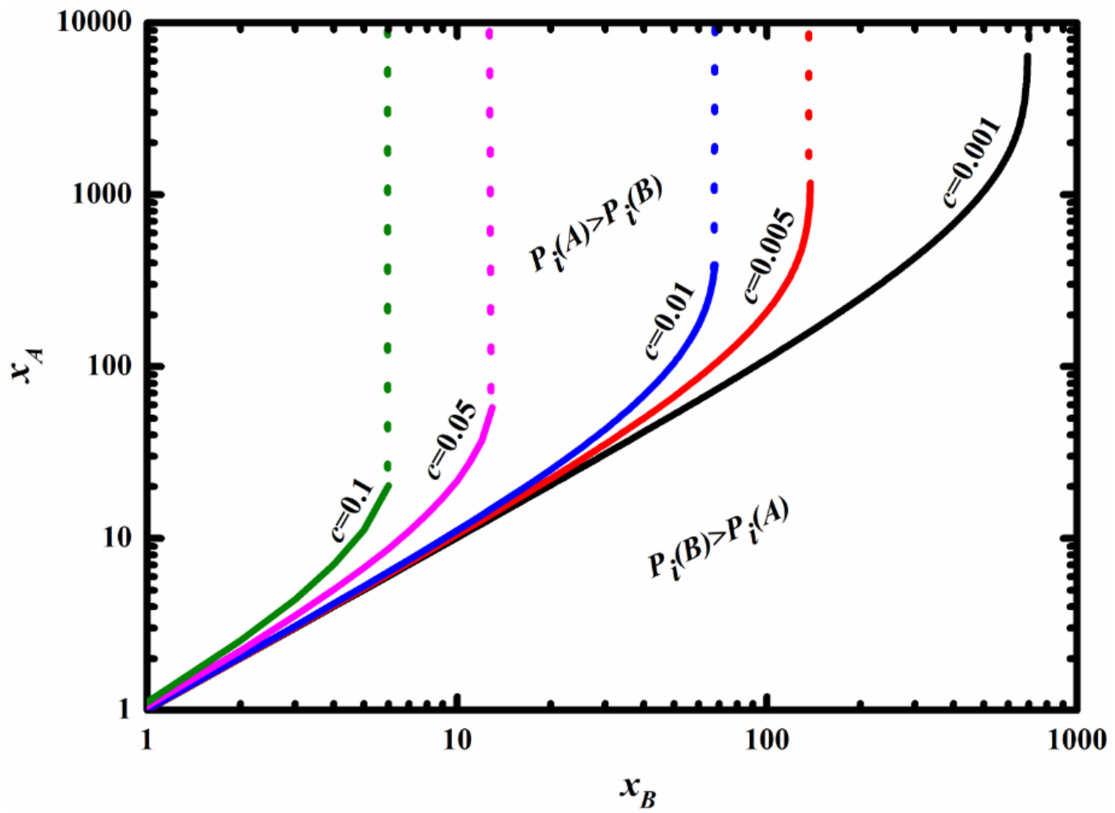


**Figure S23.** Interactive effect of the number of patches occupied separately by species $A$ and $B$ ($x_A$ and $x_B$ with the log scale) directly linked to the focal empty patch on their recolonization opportunities $P_i(A)$ and $P_i(B)$. The solid line in color dividing the region of $P_i(A)$>$P_i(B)$ (top left region) and $P_i(A)$<$P_i(B)$ (bottom right region) varies with species colonization rate ($c$=0.001, 0.005, 0.01, 0.05 and 0.1). When the values of $x_B$ are larger than the thresholds (indicated by dotted lines in colors at different $c$-values, estimated by setting $x_B = -\frac{\ln 2}{\ln(1-c)}$), species $B$ will always have higher probability to occupy the focal empty patch than species $A$ regardless of $x_A$-values, i.e. $P_i(B) > P_i(A)$.
